# A Mechanistic Model for the Release of Ceramide from the CERT START Domain

**DOI:** 10.1101/2023.12.16.571968

**Authors:** Mahmoud Moqadam, Parveen Gartan, Reza Talandashti, Antonella Chiapparino, Kevin Titeca, Anne-Claude Gavin, Nathalie Reuter

**Affiliations:** Department of Chemistry, University of Bergen, Norway; Computational Biology Unit, Department of Informatics, University of Bergen, Norway; European Molecular Biology Laboratory, EMBL, Meyerhofstrasse 1, D-69117 Heidelberg, Germany; University of Geneva, Department of Cell Physiology and Metabolism, CMU Rue Michel-Servet 1 1211 Genève 4, Switzerland

**Keywords:** CERT, STARD11, START domain, ceramide, molecular simulations, free energy

## Abstract

Ceramide transfer protein CERT is the mediator of non-vesicular transfer of ceramide from ER to Golgi. In CERT, START is the domain responsible for the binding and transport of ceramide. A wealth of structural data has revealed a helix-grip fold surrounding a large hydrophobic holding the ceramide. Yet little is known about the mechanisms by which START releases the ceramide through the polar region and into the packed environment of cellular membranes. As such events do not lend themselves easily to experimental investigations we used multiple unbiased microsecond-long molecular simulations. We propose a membrane-assisted mechanism in which the passage of the ceramide acyl chains is facilitated by the intercalation of a single phosphatidylcholine lipid in the cavity, practically greasing the ceramide way out. We verify using experimental lipidomics data that CERT forms stable complexes with phosphatidylcholine lipids, in addition to ceramide, thus providing a validation for the proposed computational model.

## Introduction

Ceramide (Cer) is the main precursor for the synthesis of signaling and complex sphingolipids present in mammalian membranes. Cer is synthesized in the endoplasmic reticulum (ER) and transported to the trans-Golgi region for further conversion to sphingomyelin for example (*1*). The ceramide transfer protein, known as CERT or STARD11, has been identified as the mediator of non-vesicular transfer of Cer from ER to Golgi at membrane contact sites. CERT consists of two domains linked by a disordered middle region (MR) (*2*, *3*): a N-terminal pleckstrin homology (PH) domain (*4*) associating with the Golgi apparatus and a C-terminal steroidogenic acute regulatory protein (StAR) related lipid transfer domain, named START in CERT (*5*) and responsible for the binding and transport of ceramide. The MR contains a serine-repeated motif (SRM) and a diphenylalanine in an acidic tract (FFAT) motif associating with ER-resident membrane proteins (Figure 1A and 1B). Regulation of CERT activity depends on the phosphorylation in the SRM and FFAT motifs. The former reduces phosphatidylinositol-4-phosphate (PI(4)P) binding and down-regulates the activity of CERT and the latter facilitates the VAP binding of the protein, leading to an enhanced ceramide transport to the Golgi (*6*). In addition, the isolated START and PH domains physically interact, suggesting the existence of an intramolecular regulation mechanism (*6*). While all the domains and motifs are required for the full activity of CERT, the START domain alone showed substantial activity for ceramide extraction and transport (*7*). Deletion of the START domain completely revokes the transfer activity, whereas mutants in which the PH or the MR domains were deleted retained this activity. The START domain is thought to bind the donor membrane via binding loops, to take up a ceramide into a hydrophobic lipid-binding cavity, and to transfer it to the acceptor membrane (*5*).

**Fig 1.**
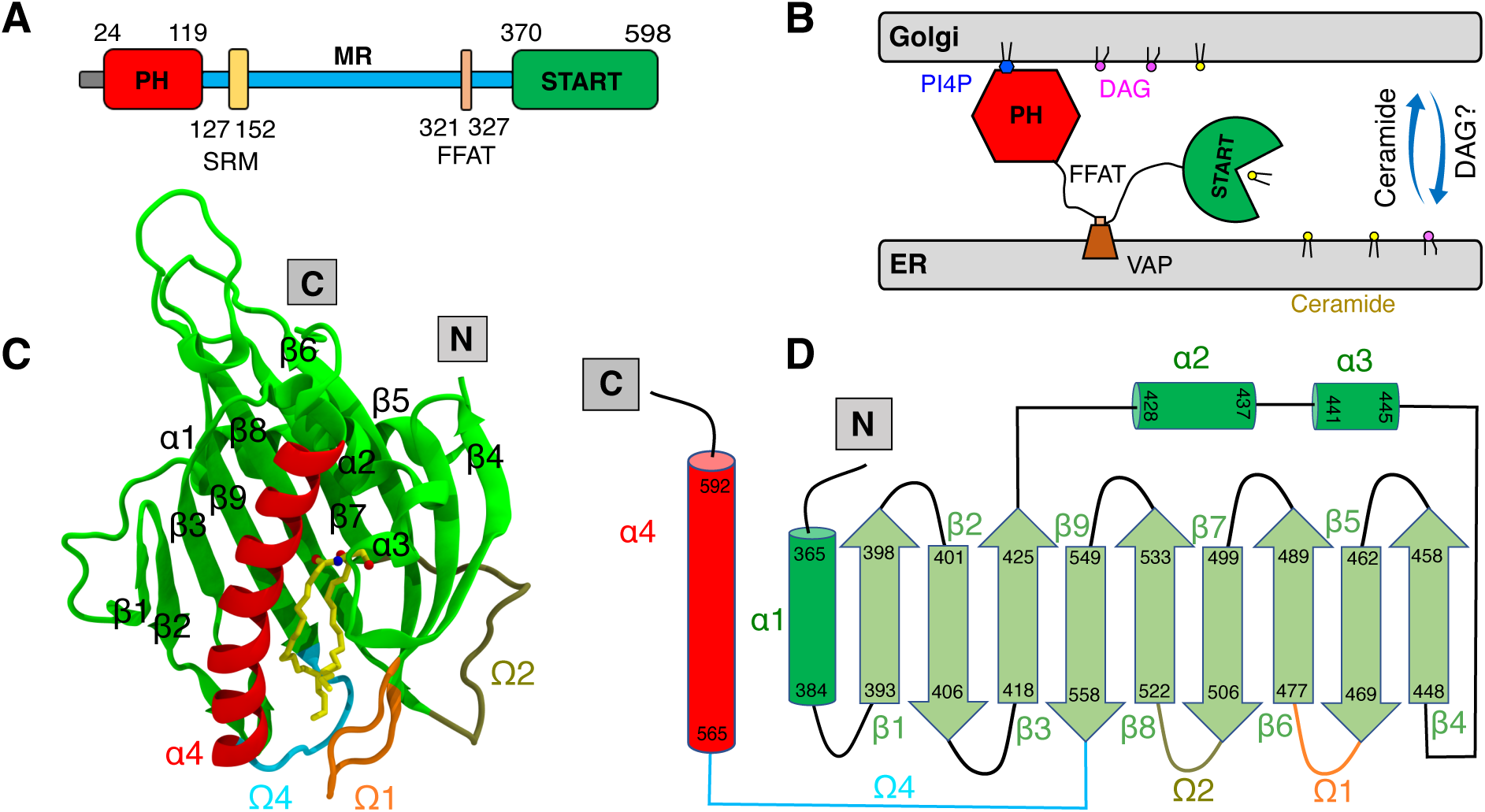
A model of CERT-mediated trafficking of ceramide. (**A**) Domains and motifs in CERT. The PH domain, START domain and sequence of the SRM and FFAT motif are depicted in colored objects, and the MR is shown as a blue bar. (**B**) Ceramide transfer from the ER to the trans-Golgi membrane. (**C**) Structure of the START domain in complex with ceramide. (**D**) Topology diagram of START domain.

CERT belongs to the StART family of the StARkin superfamily, the largest group of lipid transfer proteins (LTPs). LTPs cover a variety of folds that have as common trait a hydrophobic lipid-binding cavity shielding their cargo from the aqueous environment (*8*, *9*). The structure of proteins in the StARkin superfamily, which also includes phosphatidylinositol transfer proteins (PITPs) (*10*, *11*), proline rich EVH1 ligand 1 (PRELI) and the yeast unprocessed (Ups) (*12–14*) proteins consists of an arrangement of a beta-sheet and helices forming the hydrophobic cavity. Access to the hydrophobic cavity is thought to be controlled by at least one loop acting as a gate, but the molecular mechanism involved has not been elucidated. In CERT START domain this loop is called the Ω1 loop (exchange loop in PITP, Ω in PRELI and α2 in Ups1). The 27 available X-ray structures of START, in its apo form and bound to ceramide or various other ligands and inhibitors, show a cavity formed by residues of the α4 helix and Ω4 loop on one side (left on Figure 1C), and amino acids from the α3 helix and the Ω1 loop on the other side (Figure 1C, right). The polar headgroup of Cer is accommodated at the far end of the hydrophobic cavity forming hydrogen bond network with R442, E446, Q467, Y482, N504 and Y553 and the unsaturated sphingosine and saturated fatty acid tails are surrounded by the hydrophobic wall of the cavity, whose size and shape dictate the length limit for cognate ceramides (*15*). Kudo *et al* suggested that two exposed tryptophan residues, W473 in the Ω1 loop and W562 in the Ω4 loop (β9-α4 loop, Figure 1C and 1D), might dictate the orientation of START when bound to membranes (*5*). They showed that a W473A/W562A double mutant reduces membrane affinity with almost no ceramide extraction or transfer activities in cell-free assay systems. This mutant of CERT also shows no ER-to-Golgi trafficking of ceramide in semi-intact cells. However, the W473A/W562A CERT mutant retains the ability to localize at the Golgi region, consistent with the idea that CERT can target the Golgi apparatus by recognizing PI4P via its PH domain (*7*, *16*). Structures of START with ceramide-analog inhibitors show a large displacement of the W473 side chain which moves inside the cavity (*15*), while it is exposed to the outside in the apo-form and ceramide-bound complexes.

While Kudo et al suggested that the α3 helix (Figure 1C) and the Ω1 loop of START might function as a gate to the lipid-binding cavity (*5*), none of the 27 START structures shows an open gate, or any notable conformational differences in the region of α3 and Ω1 or in any other regions. As a consequence, the mechanisms controlling the operation of the START gate and the incorporation (or release) of Cer into (or from) the hydrophobic cavity remain poorly understood. Molecular dynamics (MD) simulations have the potential to inform about conformational dynamics of membrane-bound proteins at atomic resolution (*13*, *17–23*), and it has also shed light on uptake/release mechanisms for other members of the StARkin superfamily. Coarse-grained molecular simulations of PRELID1-TRIAP1 and PRELID3b-TRIAP1 suggest an opening of the Ω loop (Ω1 in CERT START) following insertion of the α3 helix (equivalent to α4 in CERT START) in the bilayer, and a subsequent two-step release of the bound lipid (*13*). In the equivalent protein in yeast, Ups1/Mdm35, atomistic simulations and structural data indicate that the L2 loop (equivalent to helix α3 in CERT START) also plays an important role in addition to the α2 loop (Ω1 in CERT START) and the C-terminus helix α3 (α4 in CERT START) (*24*). Interestingly, different orientations are predicted for the membrane-bound forms of PRELID1-TRIAP1 which has its long C-terminus helix nearly perpendicular to the membrane, and PRELID3b-TRIAP1 and Ups1/Mdm35 where the C-terminus helix is seen lying down on the membrane. PITPα is also predicted from all-atom molecular dynamic simulations to adopt this position (*11*), whereas for STARD4 the orientation is dependent on the membrane composition and in particular on the presence of phosphatidylinositol-phosphates (*25*, *26*). None of the reported studies using atomistic simulations did address the mechanism for lipid release and so the mechanism for it remains unknown.

Despite the wealth of structural data on the apo and holo CERT START domain with various ligands the conformational changes associated with Cer release remain elusive, and so does the mechanism by which Cer is transferred from the hydrophobic cavity, through the polar membrane interface and into the acceptor membrane. To shed light on these mechanisms we used extensive atomistic molecular simulations of the binding of the apo and holo START on complex lipid bilayers mimicking the ER and Golgi membranes, respectively. The simulations performed for each protein-membrane system were extended well beyond the membrane binding events and until two microseconds each to investigate the interplay between the protein, its cargo and the membrane lipids. The trajectories were thus analyzed with a focus on information pertinent to the opening/closing of the cavity, protein-lipid interactions and the behavior of ceramide. We observed a series of diffusive and rare events which favor the release of Cer from START domain and propose a model where these events combined would lead to full release of Cer from the holo START domain to a Golgi-like lipid bilayer.

## Results

### Atomistic molecular simulations of START domain binding to ER and Golgi membrane models

We performed molecular dynamics simulations of the apo and holo forms of START in the presence of all-atom bilayers mimicking the composition of the ER- and Golgi membranes. These bilayers are hereafter named ER and Golgi bilayers and their lipid compositions are depicted in Figure 2A. The main difference between the two is the presence of diacylglycerol (DAG) and phosphatidic acid (PA) in the Golgi-like bilayer, whereas phosphatidylcholine (PC), phosphatidylethanolamine (PE), phosphatidylinositol (PI), phosphatidylserine (PS) and cholesterol (CHL) are present in both; 1-palmitoyl-2-oleoyl (PO) chains were used for all lipids. We also used a simpler neutral bilayer consisting only of PC, PE, CHL and Cer. In what follows the simulations of the apo form on the ER-like bilayer is named apo-ER, and the simulations of the holo form on the Golgi-like bilayer is named holo-Golgi.

**Fig 2.**
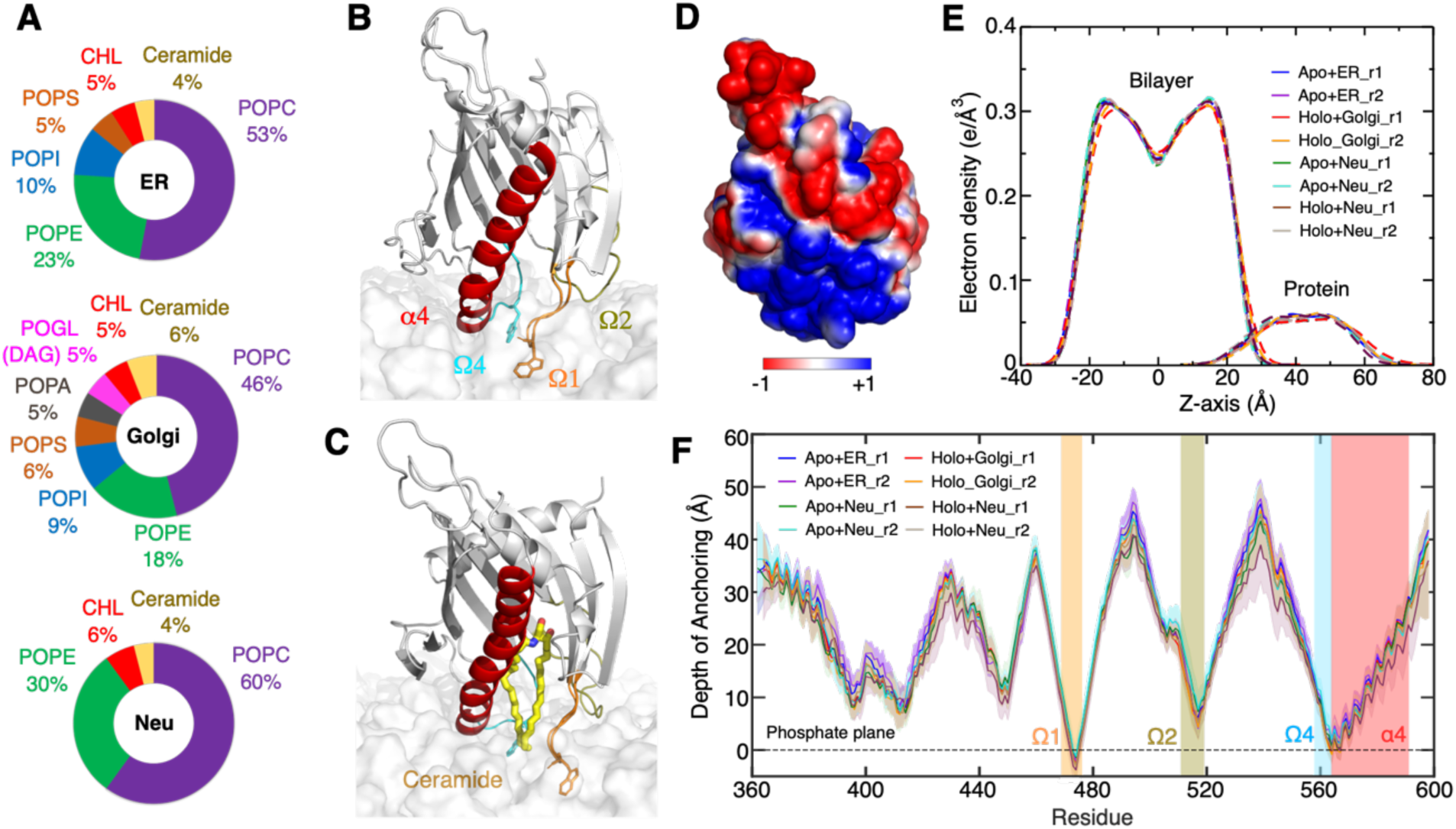
Binding mode of apo and holo START on ER- and Golgi-like bilayers from molecular dynamics simulation. (**A**) Lipid compositions in ER-, Golgi-like and neutral bilayers. (**B and C**) MD snapshots of the bilayer-bound apo and holo START on ER- and Golgi-like bilayers, respectively. (**D**) Electrostatic surface potential of apo START mapped on the molecular surface (negative: red, positive: blue) (**E**) Electron density profiles of the bilayers and protein from MD simulations including both replicas (r1 and r2) in each case, calculated for the last 500 ns of the simulations. (**F**) Depth of insertion of each START amino acids into the bilayers for both replicas (r1 and r2) in each case, calculated from the time START domain binds to the bilayer to the end of the simulations (*See Materials and Methods*).

The protein structure was initially positioned slightly above the bilayers and oriented with the Ω loops facing the bilayer. Each simulation was run for a total of 2 μs and replicated once (Table S1). Apo and holo START domains were bound spontaneously within 300 to 500 ns onto the ER bilayer and within 60 or 500 ns on the Golgi bilayer depending on the replicate. The bound orientations were close to the starting ones and are shown in Figure 2B and 2C. The structures remained close to the X-ray structures except for the N-terminal region which was highly mobile. In the full CERT protein, this region links to the FFAT motif and further to the MR domain and the absence of this domain in the simulations might explain the high mobility observed. Besides that, we observed changes in root-mean-square deviation (RMSD) (Figure S1) following the binding of START to the bilayers and corresponding to gate opening events that are described in detail in the next sections.

### The Ω1 and Ω4 loops anchor START domain to ER- and Golgi-like bilayers

To characterize the orientation and the depth of insertion of START on the bilayers we calculated the electron density of protein and lipids along the direction perpendicular (normal) to the membrane plane (Figure 2E), the angle between helix α4 and the membrane normal (Figure S2), the position of the START center of mass with respect to the bilayers mid-plane (Figure S3) and the distance between the alpha carbon of each amino acid and the average plane of the phosphorus atoms in the upper leaflet (Figure 2F).

The insertion depth and orientation of apo START on the ER bilayer is similar to that of holo START on the Golgi bilayer. There are no notable changes in the tilt angle upon binding of START to the bilayers, and the angle remains around its average value of 40°±15° for the ER and Golgi bilayers (Figure S2). For both types of membranes, START (apo on ER and holo on Golgi) is anchored rather superficially to the bilayers (Figure 2E) through the C-terminus α4 helix, the Ω1, Ω2 and Ω4 loops (Figure 2F). Only a few residues are inserted under the average plane of the phosphorus atoms (Figure 2F). This is also reflected in the inventory of interactions between START amino acids and the bilayer lipids (Table S2). Helix α4, Ω1 and Ω4 mediate hydrophobic contacts with multiple lipid tails mostly through the same amino acids in both bilayers; W473 (Ω1) and W562 (Ω4) and a few hydrophobic amino acids V472 and P474 in Ω1, V562 and P564 in Ω4, V571 in α4. Helix α4, the Ω1, and Ω4 loops also engage in several long-lasting hydrogen bonds with the lipid head-groups, mostly with the phosphate groups. Six basic amino acids are involved: R471 in Ω1, R478 in β6 (right after Ω1), R569, K573, R574 and K578 in Ω4. In addition, we observe hydrogen bonds with R517 in the Ω2 loop. Unlike what we have observed for other proteins (*27*), there are no long-lasting cation-π interactions between aromatics amino acids and choline headgroups (*28*). The two exposed tryptophane residues, W473 and W562, engage in hydrophobic contacts with the fatty acid tails and hydrogen bonds with the phosphate and glycerol groups of the membrane lipids. W473 is inserted deep in the bilayer forming hydrogen bonds with the lipid glycerol group, while W562 mostly forms hydrogen bonds with the lipid phosphate groups (Table S3).

The electrostatics surface potential of START domain (Figure 2D) shows a positive region at the membrane-binding site suggesting an electrostatic recognition of the ER and Golgi bilayers, which both contain negatively charged lipids. We evaluated the influence of the surface charge of the bilayers on protein binding by performing a control simulation with an ER bilayer stripped from its negatively charged lipids (Neu on Figure 2A). There was no difference in the depth of insertion (Figure 2E and 2F) or the orientation (Figure S2) of START on the neutral Neu bilayer compared to the ER and Golgi bilayers, ruling out a dominating electrostatic contribution in START membrane binding. Overall, our data suggests that the START domain has general affinities with membranes and that this domain alone is not sufficient to target specific lipids or bilayers. This function is probably mediated by the PH domain, the SRM and/or FFAT motifs of CERT.

### The opening of Ω1 and Ω4 loops triggers snorkeling of lipid tails under the hydrophobic cavity of START domain

Visual inspections of simulations of START (apo and holo) on the ER and Golgi bilayers revealed an opening of the gate to the ceramide binding pocket through displacements of the N-terminal end of helix α4 and of loops Ω1 and Ω4. The change was not observed in the second replica of the START-ER simulation. The conformational change is illustrated on Figure 3A with a snapshot taken at t≈800 ns of the simulation of the apo form of START on the ER bilayer. Shortly after the binding of START onto the bilayers, α4 and the Ω1 and Ω4 loops undergo a large structural shift leading, among other things, to the disruption of interactions between the two loops (Figure S5). The gate opening is quantified on Figure 3B by time series of the distances between P564 (α4) and W473 (Ω4) on the one hand, and W562 (Ω1) and S476 (Ω4) on the other hand (see Figure S4). Furthermore, Figures 3C and 3D show the distributions of the closed and opened states of START domain on the ER and Golgi bilayers. The average distances of the α4 helix and Ω4 loop to the Golgi bilayer are 9.5 Å and 15 Å for the closed and open states, respectively. Upon opening of the gate, the side chains of W473 and W562 shift toward the outside of the cavity resulting in a fully open state (Figure 3A).

**Fig 3.**
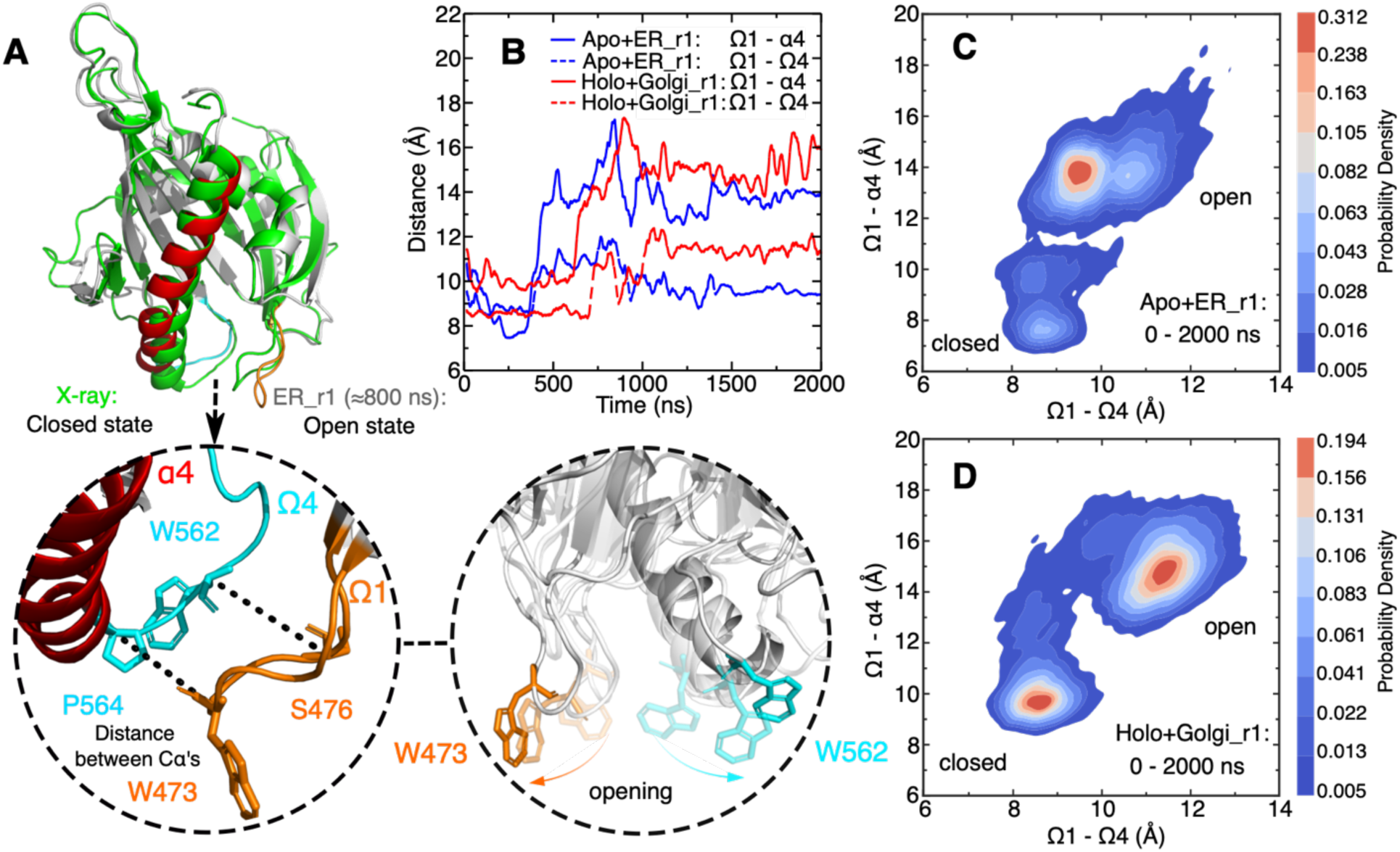
Gate opening through displacements of Ω1, Ω4 and α4. (**A**) Superimposition of the open state in grey cartoon (α4 in red, Ω1 in orange, Ω4 in cyan, and MD snapshot at ≈800 ns of apo+ER_r1 simulation) with the closed state in green cartoon (X-ray structure, PDB ID 2e3m) and close-ups of the open state and of the orientation of the tryptophane W473 and W562 side chains along a simulation. Hydrogens are not shown for the sake of clarity. (**B**) Time series of the Ω1-α4 (plain lines) and Ω1-Ω4 (dotted lines) distances in simulations of START with the ER (blue) and Golgi (red) bilayers (See Figure S4 for the replicas). (**C and D**) Density distributions of the closed and open states of START simulated with the ER bilayer (C), the Golgi bilayer (D).

We also performed a 1 μs-long simulation of the START apo form in water and replicated it twice. The analysis of these simulations, specifically the time series of the START domain opening distances (Figure S6 and S7), demonstrates the occurrence of opening events similar to those observed in membrane-bound START domain. However, these events have a short duration of only 100 ps. This indicates a role of the membrane lipids in stabilizing an opened form of the gate, rather than triggering it.

START opening exposes the hydrophobic lipid-binding pocket to the hydrophobic environment of the bilayer. In the simulation of the apo START on the ER bilayer, and of the holo form on the Golgi and neutral bilayers, the opening of the gate is concomitant to a change of the neighboring lipid tails that tend to snorkel in the space just under the opened hydrophobic cavity. This is illustrated by the simulation snapshots on Figure S8 and quantified by monitoring the distance between the last carbon atoms of the lipid tails and the average position of the phosphate groups, projected onto the normal to the bilayer (with respect to Z axis) (Figure S9). From the gate opening event in the apo-ER_r1 simulation, and for as long as it is opened, we observe between 3 and 4 lipids positioned under the gate and whose tail deviates from the orientation of other lipids. For the sake of comparison, the average number of lipids undergoing such conformational changes in the whole opposite leaflet (ie in the absence of protein) is between 1 and 2 (Figure S10). Interestingly, in the holo-Golgi simulation, these events led to the bound ceramide to extend its tails towards the bilayer, increasing the number of contacts between ceramide and lipid tails from 0 to 10-16 before and after gate opening in the first simulation, and from 0 to 20-24 contacts in the replicate (Figure S11).

### POPC tail insertion triggers release of the ceramide

Besides the frequent local molecular rearrangements of proteins and the lipid tails in the bilayers described above, we observed the engagement of a POPC (palmitoyl-oleoyl-PC) lipid in the hydrophobic cavity. That POPC lipid inserts one tail into the hydrophobic cavity in one replicate of each of the apo-ER, apo-neutral, holo-neutral, and holo-Golgi simulations (see supplementary text S1 for other replicates). This phenomenon is illustrated by the snapshots in Figures 4 and S12. The orientation of the inserted tail is quantified by its angle with the membrane normal (Figures 4B, 4D, S12B, and S12D). In each of the four simulations a POPC phosphate group is locked in the gap between the opened Ω1 and Ω4 loops by a salt bridge with R478 and hydrogen bonds with the S476 (Figure S13). One tail of that POPC engages in hydrophobic contacts with exposed amino acids at the entrance of the cavity and shifts toward the hydrophobic core of the cavity to finally insert into it (Figures 4A, 4C, S12A, and S12C). The tail angle then changes from high values when the tail is in the bilayer (above 150 degrees) to low values characteristic of the insertion in the cavity (45 degrees and below). There are slight differences in the four simulations. In three of the simulations (apo-ER, apo-Neu, holo-Neu) the unsaturated tail is inserted, while the saturated fatty acyl chain stays in the membrane. In the fourth simulation (holo-Golgi), it is the saturated tail that moves towards the gate, but it adopts a somewhat different position than the unsaturated tail in the other three simulations. Indeed, it interacts mostly with the Ω1 loop at the entrance of the cavity (Figure S12C), unlike in the other three cases where it interacts with residues located deep in the cavity (Figures 4A, 4C, and S12A). The lifetime of the START-POPC complex is short in the holo-Golgi (40 ns) and apo-Neu (70 ns) simulations, but longer in the apo-ER (500 ns) and holo-Neu (750ns) simulations (see Movie S1, S2A, S2B, and S3).

**Fig 4.**
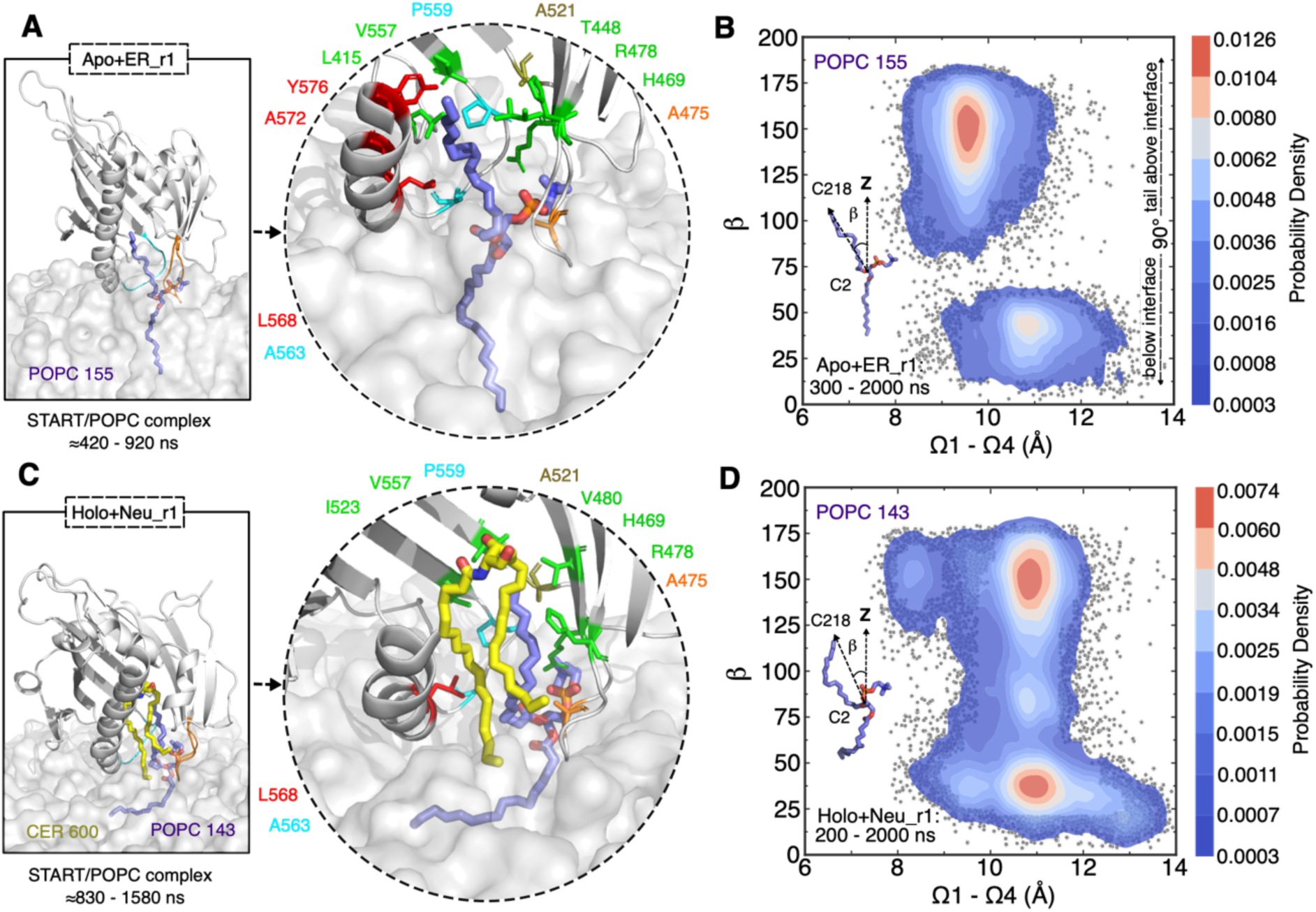
POPC (1-palmitoyl-2-oleoyl-phosphatidylcholine) rearrangement induced by START domain in opened state detected by MD simulation. (**A**) Close-up view of the binding conformation of POPC within apo START domain, and the amino acids in the cavity involved in hydrophobic contacts with the POPC tail in the ER bilayer. (**B**) Density distribution of the POPC tail angle with respect to membrane normal in the ER bilayer is presented. (**C**) Close-up view of the binding conformation of POPC within START-Cer complex, and the amino acids in the cavity involved in hydrophobic contacts with the POPC tail in the neutral bilayer. (**D**) Density distribution of the POPC tail angle with respect to membrane normal in the neutral bilayer is presented. The times at which the POPC tail inserts into and exits the cavity are given below the snapshots. Ceramide colored yellow, POPC colored purple, Ω1 colored orange and Ω4 colored cyan. START domain is shown as cartoon and colored grey. Membrane is shown as grey transparent surface. The gray dots represent all the sampled tilt angles.

Following the insertion of a POPC tail into the cavity in one of our simulations of the START-Cer complex (holo-Neu, Figure 4C), we observe a temporary release of ceramide from the cavity and into the lipid bilayer (Figure 5A-F). The POPC tail intercalates between the protein and the ceramide. The time series of the hydrophobic contacts before and after the tail insertion shows a disruption of the protein-ceramide hydrophobic contacts (with R478, V480, A521, I523, V557, and P559) and the formation of contacts between these residues and POPC (Figure 5H-I). The hydrogen bonds between the ceramide headgroup and its binding site (Y482, N504, and Y553) then break sequentially (Figure 5G). This step is followed by the release of ceramide which inserts between other lipids in the bilayer (Figure 5E and S11). It is followed by the exit of the POPC lipid tail from the cavity (t≈1580 ns). Subsequently the ceramide moves back up to the cavity (Figure 5F) but its head group fails to re-establish a stable hydrogen bond network with Y482, N504, and Y553. Instead, the ceramide slides back and forth along the long axis of the cavity for the remaining simulation time (see Movie S2C).

**Fig 5.**
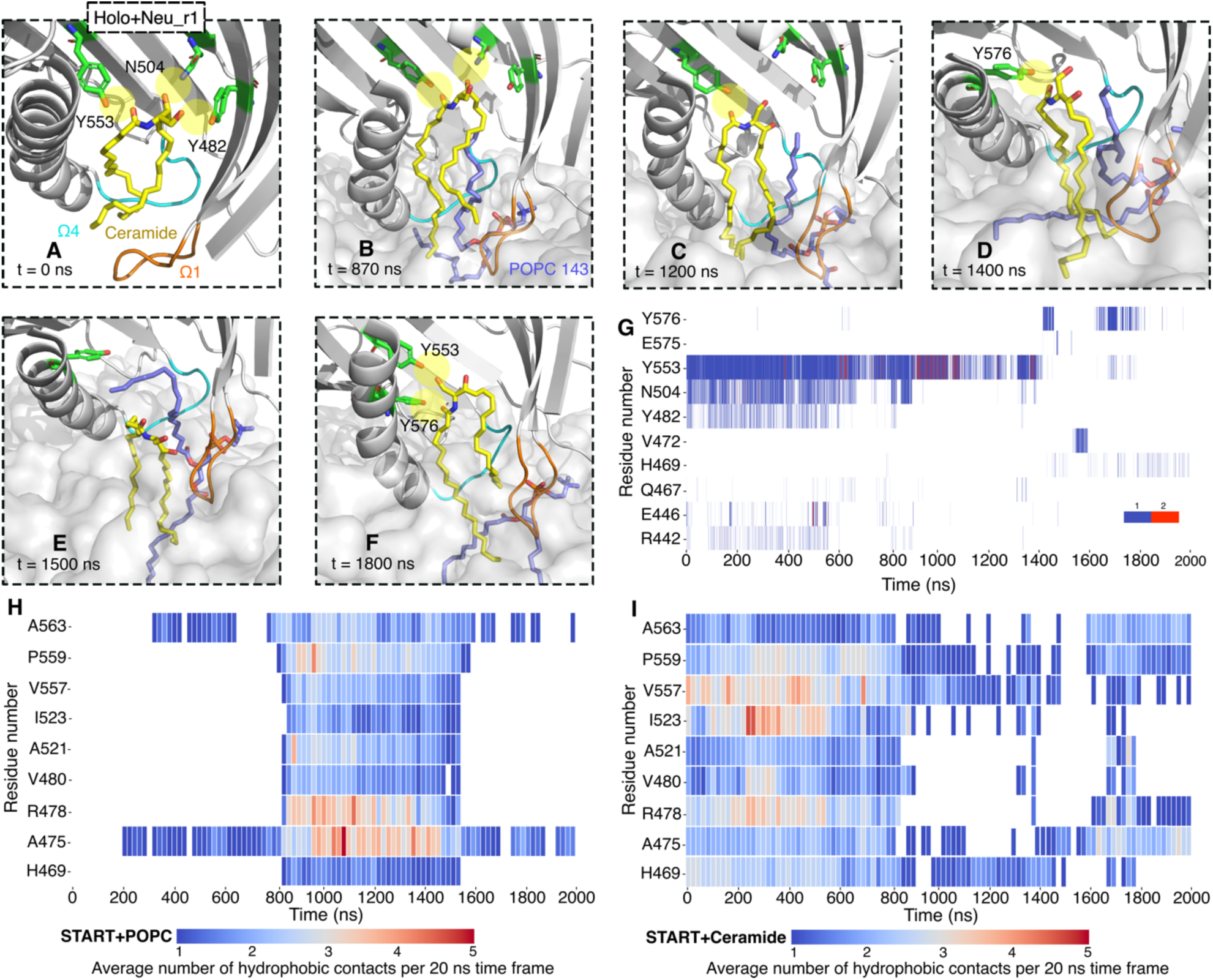
Snapshots along 2 μs simulations of holo START on the neutral bilayer. (**A**) Hydrogen bond network between the ceramide and cavity residues (green licorice, yellow highlight for hydrogen bonds) before binding to the bilayer. (**B and D**) Hydrogen bonds break one by one, and the Cer is displaced from the binding site, forming a hydrogen bond with Y576 of the C-terminus α4 as the Cer moves toward the bilayer (see Figure S14 and Table S4 for other replicates). (**E**) The POPC143 tail disrupts the Cer-Y576 hydrogen bond. (**F**) Cer forms hydrogen bonds again with Y553 and Y576 as the POPC tail leaves the cavity. (**G**) Time series for hydrogen bond network between ceramide and START domain. Color bar indicates the number of hydrogen bonds in each frame. (**H and I**) Time series of the average number of hydrophobic contacts for selected residues, calculated in 20 ns time windows, throughout the 2μ simulations: (H) POPC143 with the protein and (I) ceramide with the protein. (see Figure S15 for all the hydrophobic contacts). Ceramide colored yellow, POPC colored purple, Ω1 colored orange and Ω4 colored cyan. START domain is shown as cartoon and colored grey. Membrane is shown as grey transparent surface.

We observe other lipids between Ω1 and Ω4 during our simulations, such as POPE (palmitoyl-oleoyl-PE) or POPS (palmitoyl-oleoyl-PS) but no other lipid than PC do insert a tail in the cavity. Like POPC, the PE and PS lipids form hydrogen bonds with neighboring amino acids (S476, R478, W562) but unlike POPC, their headgroups (ethanolamine and serine) are engaged in long-lasting interactions. These interactions are likely to explain the limited degrees of freedom of the bound PE and PS lipids that we observe, and the apparent restricted mobility, which hinders the insertion of their tails in the hydrophobic cavity.

Our observations from the holo-neutral and holo-Golgi bilayers suggest that the ceramide release is triggered by a succession of events: (1) the hydrophobic cavity is stabilized in its open form by the lipid bilayer, (2) the ceramide exposed tails are in contact with bilayer lipids, (3) bilayer lipid tails snorkel around the open cavity, and (4) a POPC tail is inserted deep into the cavity (such as in the holo-neutral simulation). Steps (1)-(3) occur in every simulation replicate while step (4) is observed in half of the replicates suggesting that it might be a diffusive rare event (other replicates are reported in Supplementary Text S1).

### Verifying that POPC intercalation between ceramide and START disrupts protein-ceramide interactions and triggers cargo release

We first evaluate the effect of the loss of START-Cer interactions on the stability of the complex. The head group of ceramide forms hydrogen bonds with the START domain (Figure 5B and 5G) and the ceramide tail forms hydrophobic contacts with several amino acids (Figure 5I). The simulation results show a gradual disappearance of these interactions, partly under the influence of the inserted POPC tail but also prior to it. We here evaluate the contribution of a selection of these interactions to the START-Cer affinity to verify that their disappearance is likely to modify the stability of Cer in the START cavity and favor its release. First, we calculated the contributions to the START-Cer binding affinity of the Y553 and N504 hydrogen bonds, and of the V480 hydrophobic contact to Cer. This was done by calculating the energy change associated with the Y553F, N504A and V480A substitutions, in the presence and absence of ceramide (see Methods section and Tables S5 and S6). We find that the cost of the Y553F and N504A substitutions is 0.8 and 1.1 kcal/mol respectively, confirming the favorable contribution of Y553 and N504 to the START-Cer affinity. The V480A substitution is unfavorable (1.2 kcal/mol) indicating that V480 contributes favorably to the START-Cer interaction. The presence of the POPC tail reduces this contribution to a negligible value (0.2 kcal/mol). Overall, these data indicate that the three amino acids contribute favorably by about 1 kcal/mol to the START-Cer binding affinity and so their loss is likely to weaken the affinity of Cer for START. As expected, the insertion of the POPC tail in the cavity removes the favorable contribution of the V480 to Cer binding.

As another verification of our proposed model of POPC interfering with the START-Cer interactions, we built a slightly modified holo-Golgi system. We positioned the POPC tail similarly to that which led to the temporary Cer release in the holo-neutral simulation and shown on Figures 5A-E (*See Material and Methods for information about model building*). We then subjected the model to a 2 μs MD simulation in the presence of the Golgi-like bilayer and replicated the calculation once. In both replicates, the ceramide fully leaves the cavity and enters the bilayer where it mixes with other lipids. Figure 6 shows the initial structure (open form, Figure 6A) anchored to the bilayer (Figure 6B). Within 100 ns the hydrocarbon tails of the ceramide engage in interaction with the lipid bilayers, and within 1 μs of the simulation (500 ns in the second replicate) the ceramide is fully released into the bilayer (Figure 6C). Notably, we also observe the uptake of a POGL lipid in one of the replicates (Figures 6D-F) but it is released within 200 ns into the bilayer (see Movie S4). These simulations confirm that when inserted in a position which weakens the Cer-START interactions quantified above, the POPC lipid will trigger the release of the ceramide cargo on the Golgi-like bilayer.

**Fig 6.**
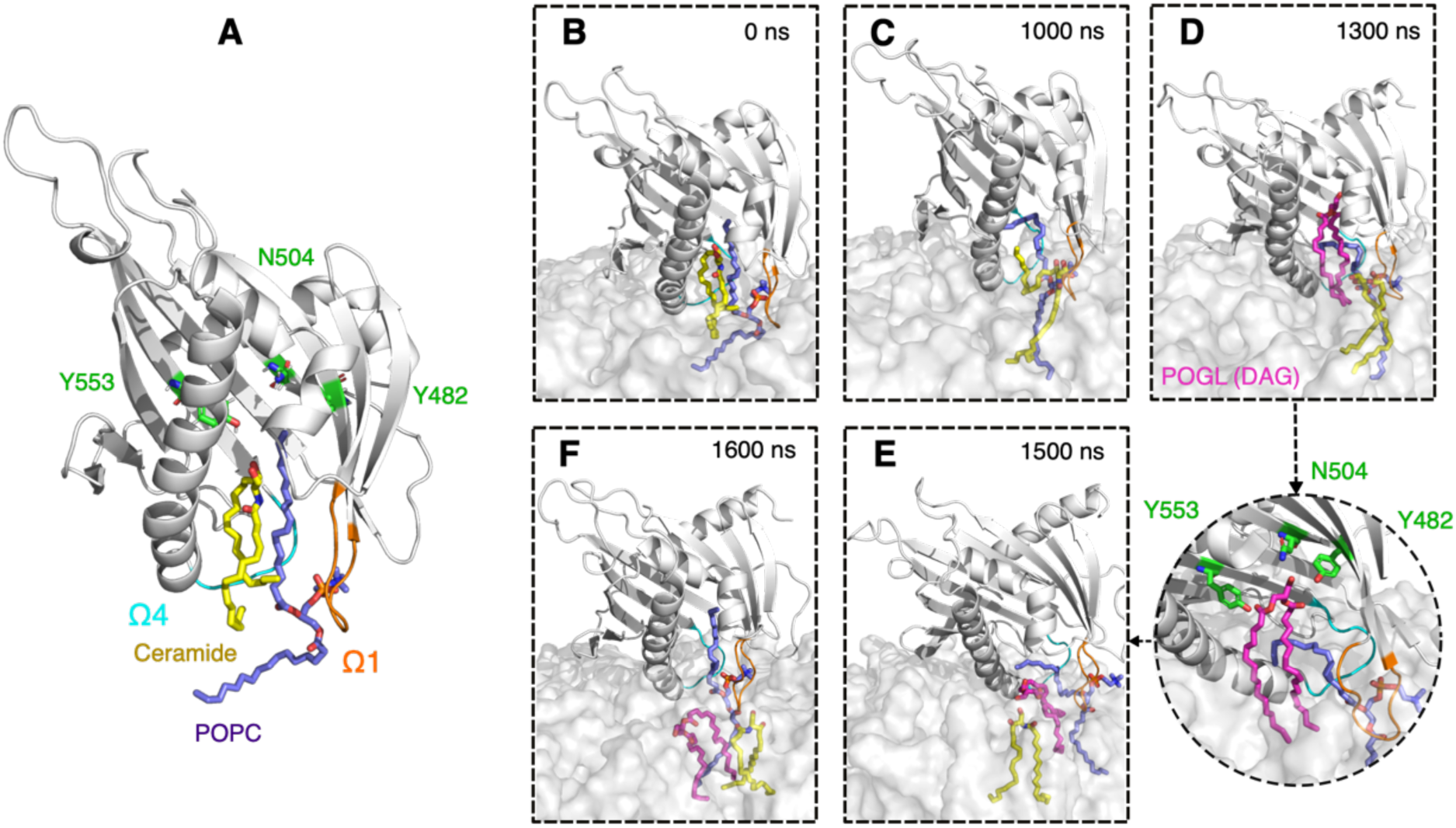
Stages of membrane binding and ceramide release in the model system (Golgi bilayer). (**A**) Built initial complex of START, ceramide, and POPC. (**B and C**) the ceramide release to the bilayer. (**D-F**) The diacylglycerol lipid (POGL) uptake (and release) from (to) the bilayer. Bound ceramide molecule colored yellow, POPC molecule colored purple, residues in the binding site colored in green. POGL is shown in magenta and bound in the cavity. Ω1 loop colored orange, Ω4 loop colored cyan and START domain colored grey. Membrane is shown as grey transparent surface.

### CERT forms complexes with ceramide and phosphatidylcholine in human HEK293 cells

The *in silico* models suggest that CERT-mediated ceramide transport requires direct interaction with phosphatidylcholine in the membrane, and temporarily at least partial uptake in its hydrophobic cavity. We set out to validate this experimentally by integrating and reanalyzing data from our recent systematic biochemical analysis of LTP cargoes. We over-expressed CERT in HEK293 cells and, after biochemical purification, characterized CERT-associated lipids by LC-MS/MS-based lipidomics (Figure 7 and Table S7. We have identified 32- to 36-carbon ceramide species, known CERT cargoes, as well as PC (34:1) and PC (32:1) forming stable complexes with CERT (Table S8). This is the first observation of the association of PC with CERT from a real cellular context. Phosphatidylcholine is one of the two substrates (with ceramide) of the sphingomyelin synthase, an enzyme in the trans-Golgi known to accept ceramide from CERT for its function. Sphingomyelin synthase transfers the phosphocholine headgroup of phosphatidylcholine to ceramide to generate sphingomyelin and diacylglycerol. This is consistent with the *in silico* observation of both PC and ceramide in the START cavity, and the dependence of the release of ceramide on the presence of PC in the target membrane, thus ensuring the presence of PC and ceramide in the same location, being the two substrates of a known downstream enzyme sphingomyelin synthase.

**Fig 7.**
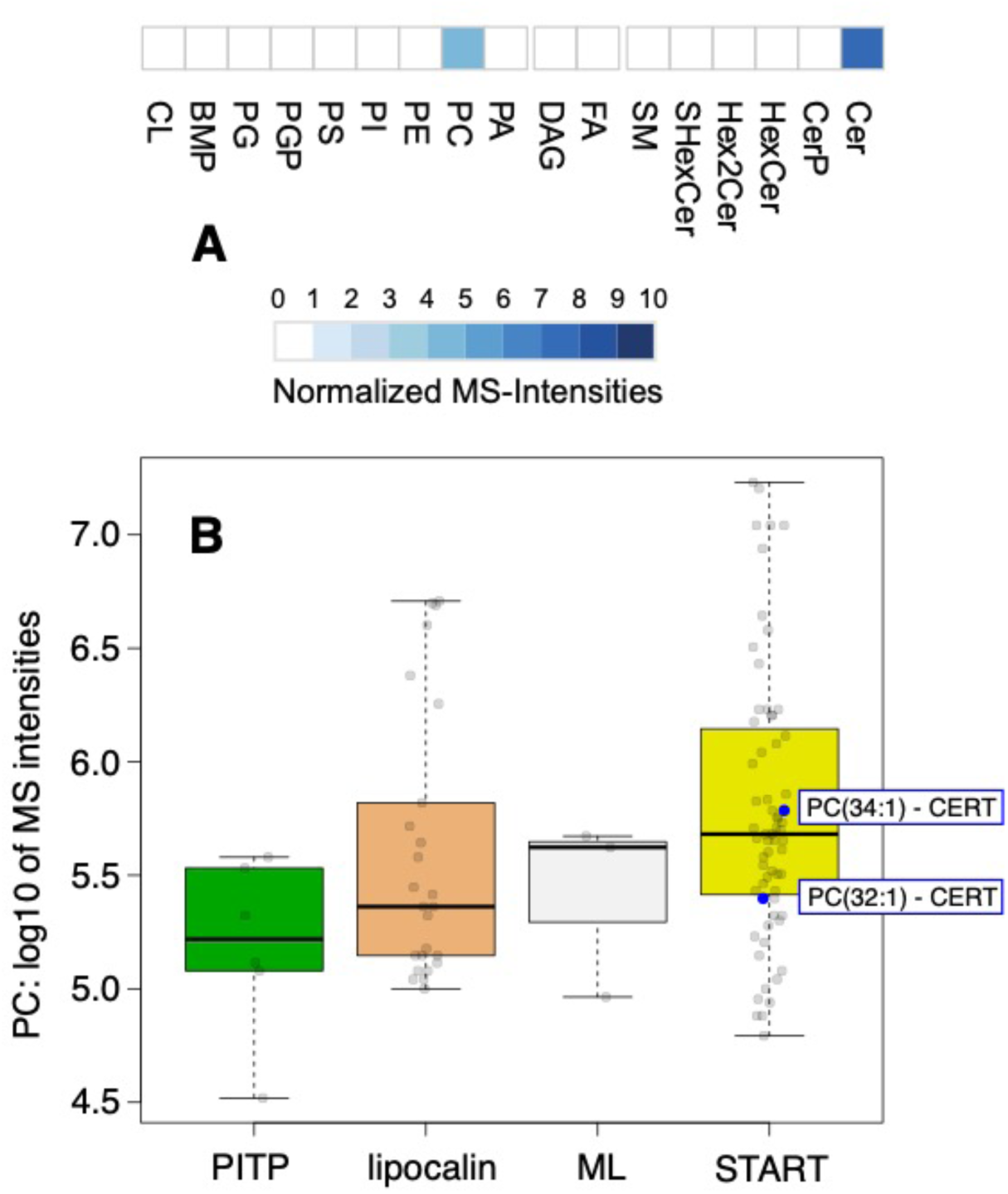
CERT expressed in HEK293 cells forms complexes with ceramide and PC. Biochemical purification and characterization of CERT-lipid complexes by LC-MS/MS-based lipidomics. (A) Quantification of CERT-associated lipids by LC-MS/MS. The heat map represents the normalized sum of the MS intensities of all lipid species for a given lipid class. (B) Comparison of the PC-binding capacities of CERT with that of other LTPs known to form complexes with PC and belonging to the PITP family (PITPNA and PITPNB), lipocalin family (LCN1), ML family (GM2A), and START family (STARD2, STARD10 and CERT).

## Discussion

Herein, we report a series of extensive unbiased MD simulations of CERT START domain on lipid bilayers whose lipid compositions mimic that of the cytoplasmic leaflets of the Golgi apparatus and the endoplasmic reticulum. Lipid transfer mechanisms rely on the concomitant occurrence of multiple diffusive and rare events and the likelihood of observing all required events concurrently within the time frame of unbiased μs-long MD simulations is low. Combining detailed analysis of the MD trajectories with free energy calculations and experimental data, we are able to propose a mechanistic model for the interfacial processes leading to ceramide release by CERT START domain at the Golgi membrane. In this model, the membrane plays a major role both in the gate opening and cargo release.

According to our simulations, the membrane binding site of the START domain consists of the Ω1 loop, the N-terminal end of α4 helix and the Ω4 loop. Hydrophobic amino acids in these three regions engage in hydrophobic contacts and in hydrogen bonds with the lipids. This is in agreement with experimental data (*5*, *15*) and in particular the reduction of membrane affinity observed for the START W473A mutant and for the double mutant W473A/W562A, and the drastic effect it has on ceramide transfer (*5*). W473 (Ω1) and W562 (Ω4) are observed engaging in long-lasting interactions with lipids in our simulations. Interestingly we do not observe differences in the membrane-binding orientation of the apo and holo forms of START domain, and the orientation is not sensitive to the lipid composition of the bilayer, indicating a lack of lipid specificity by the START domain itself. This is compatible with the fact that CERT, in addition to its START domain, contains the PH-domain and FFAT-motif which ensure targeting of the correct membranes (*29*, *4*).

Our results suggest that Ω1 and Ω4 form the gate through which the cargo enters and leaves the hydrophobic cavity. We observe a spontaneous opening of the Ω1 and Ω4 loops, accompanied by movements of the N-terminal end of helix α4, following membrane binding. Upon opening, W473 and W562 undergo a large structural shift toward the outside of the cavity thus exposing the hydrophobic pocket and the bound ceramide to lipids from the bilayer. In the absence of lipid bilayers, ie when START domain is simulated in water, we observe only transient short-lived opening events indicating a major role of the membrane in maintaining the open state. This would explain the difficulty in observing the open state with structure resolution methods where no membrane is present. The membrane-bound orientation and opening we observe are comparable to that reported for PRELID-TRIAP1 (*13*), but do not match with the hypothesis that the Ω1 loop and the α3 helix might function as a gate to the cavity (*5*). We do not observe an involvement of α3 in contacts with the bilayer lipids, or any notable structural changes that would indicate an involvement of α3 in modulating the access to the cavity. Yet, our observation agrees with the drastic effect of the W473A/W562A double mutant on ceramide transfer: W473A sits on the Ω1 loop and W562 on the Ω4 loop.

Following the opening of the gate the lipids located below the START domain undergo a reorientation of their tails which will snorkel towards the hydrophobic cavity. On the Golgi-like, ER-like or Neu bilayers, we observe a POPC lipid tail inserting into the hydrophobic cavity while the polar headgroup interacts with S476 and R478 maintaining the lipid between the Ω1 and Ω4 loops. Interestingly S476 (Ω4) is conserved in STARD2-6, and R478 (β6) is conserved in all START domains, except STARD13 (H) and STARD14 (Q). Lipid snorkeling is a known phenomenon that has been observed in both homogeneous POPC and very complex brain membranes (*30*, *31*). It is thought to be favored in heterogeneous membranes, and is correlated with increasing degrees of unsaturation in the hydrocarbon chains as well as longer tail length (*31*). In our simulations the POPC tail insertion leads to a weakening of the interactions between START and its cargo Cer, and a displacement of the cargo down towards the lipid bilayer. In two simulations, we observe a spontaneous release that can be summarized as two consecutive events: (1) the inserted tail disrupts the hydrophobic packing between the bound ceramide and the cavity and weakens the Cer-START affinity, and (2) a disruption of the hydrogen bond network between ceramide and the three amino acids S482, N504 and Y553.

Interestingly, we could not observe a full ceramide release in the presence of an inserted POPC tail in the simple POPC:POPE:CHL:Cer membrane, while it happened on the Golgi-like bilayer which in addition contains a diacylglycerol (POGL) and the negatively charged lipids POPI, POPS, and POPA. It also contains more ceramide than the ER-like and neutral bilayers. POPA and diacyglycerols (DAG) are conical lipids which can influence the phospholipid bilayer structure (*32–34*). A relevant effect is the so-called umbrella effect whereby DAG and CHL avoid the unfavourable exposure of their hydrophobic parts to water by hiding their small headgroup under neighbouring phospholipid headgroups. This increases the spacing between phospholipid headgroups and the exposure of their hydrophobic tails (*32*, *35–38*). This accessibility to lipid tails can increase the contacts between the ceramide exposed tails and lipids bilayer, which we suggested as one of the four required events for triggering the release of the ceramide. Yet, it is important to note that the amount of DAG and CHL is low (5% each) in our Golgi-like bilayers and that we can only speculate on the effect of the bilayer composition.

The transfer activity of LTPs is thought to be influenced by the lipid composition of the donor and acceptor membranes (*39–42*). Moreover it has been suggested that CERT extracts ceramide from a ceramide enriched platform (*43*). The membrane environment affects the lipid miscibility and interaction with neighboring lipids, as well as the ability of the proteins to scan the membrane surface for the target lipid (*43*). Our observations indicate that the protein itself strongly influences the local lipid packing upon binding to the bilayers. It induces changes in the organization of lipids which form a hydrophobic pool under the protein, surrounded by lipid headgroups. This interfacial hydrophobic pool provides a non-polar environment that shields the hydrophobic lipid tails from contact with water or the polar headgroups of the membrane lipids. This results in looser lipid packing and increased mobility of the lipid tails toward the open hydrophobic cavity.

Interestingly, the binding and opening mechanisms of the START domain do not appear to be influenced by lipid composition, suggesting the domain’s lipid-insensitive nature. However, the cargo release mechanism demonstrates a sensitivity to the membrane lipid composition, implying a potential role of the membrane in the release of the ceramide. It is logical to assume that other domains within CERT may be responsible for regulating membrane binding and potentially modulating the orientation of the START domain relative to the membrane. Notably, the MR domain contains distinct regions and motifs, such as the FFAT motif and phosphorylation sites, which play crucial roles in CERT function and regulation (*44*). Additionally, the interaction of the PH domain with PI4P in the Golgi membrane is essential for maintaining CERT at the Golgi surface wherein CERT releases its ceramide at a certain threshold of ceramide level (*43*). The dependence on PC in the Golgi membrane for the release of ceramide as cargo from CERT also necessitates their presence in the same location at the same time. This is particularly interesting because PC and ceramide are known substrates for sphingomyelin synthase, which is already known to be downstream of the ceramide transport by CERT. This could furthermore be consistent with the hypothesis that CERT could act as a chaperone, a notion already proposed for the CRAL-TRIO family of LTPs (*45*), potentially bringing both substrates to the same location, and coupling the transfer to their transformation into sphingomyelin. Together, it seems that the CERT binding and lipid transfer mechanisms are regulated by many factors that we are only beginning to understand.

Following the full release of ceramide, we observed POGL extraction from the bilayer. POGL structurally resembles ceramide and is produced as a byproduct in the synthesis of sphingomyelin wherein the phosphorylcholine head is transferred from phosphatidylcholine to ceramide, liberating POGL. It has been shown that the CERT START can transfer short-chain diacylglycerol (C10-DAG) *in vitro* but with a lower efficiency (only 5%) of the ceramide suggesting that CERT might mediate Golgi-to-ER trafficking of DAG in exchange for ER-to-Golgi trafficking of ceramide (*46*, *47*). In our simulation the POGL lipid does not remain in the cavity for long. This could be due to the longer acyl chain of the POGL (16:0-18:1) in our simulation (compared to C10-DAG) as well as the presence of the POPC tail in the cavity that itself facilitates the cargo release process. The impact of the presence of POPC tail in the cavity for the lipid extraction is also evident in Movie S2C (holo+Neu_r1), where the partially released Cer returns to the cavity only when the POPC tail exits. These observations might interestingly give some insight into the CERT lipid uptake from the donor membrane. The lipid tail insertion contributes to the opening of the cavity and maintains the opening in apo START easing the Cer uptake from the donor membrane while facilitates the Cer release from the holo START into the acceptor membrane.

Little is known about the atomistic-level mechanisms by which LTPs extract and release lipids from cellular membranes - or even from simpler in vitro vesicle models - because these events do not lend themselves well to experimental investigations. Molecular dynamics simulations have the potential to provide mechanistic models and generate hypotheses testable experimentally (*13*, *17–23*). Microseconds-long molecular dynamics simulations of START in the presence of lipid bilayers led us to propose a membrane-assisted model of ceramide release from CERT where (i) the membrane lipids stabilize the gate in an open form exposing the large hydrophobic cavity to the membrane interface leading to (ii) an increase of local lipid disorder and (iii) the intercalation of a single PC lipid in the cavity facilitating the passage of the cargo hydrophobic chains through the polar membrane interface, practically greasing its way out. We then set out to experimentally challenge the last step and verified that phosphatidylcholine lipids can indeed form stable complexes with CERT. Altogether our results underline the critical active role played by the membrane lipids in CERT binding and cargo release. Further investigations will be needed to evaluate a potential generalization of the proposed mechanism to other members of the StART family which, given their fold similarity may be operating in a similar fashion.

## Materials and Methods

### Systems preparation for apo-ER, holo-Golgi and holo-neutral

We retrieved X-ray structures from the Protein Data Bank (PDB) for apo CERT START (PDB id: 2e3m (*5*)), holo CERT START (PDB id: 2e3q (*5*)) domain. The apo and holo structures contain coordinates for amino acids V362-F598 and T364-F598, respectively. The two protein structures were placed slightly above the surface of the relevant lipid bilayers (ER-, Golgi-like, and neutral) built using CHARMM-GUI (*49*), explicit TIP3P water molecules (*50*), and neutralizing potassium ions. The starting orientation of the protein on the bilayers was obtained from theoretical predictions from the Orientation of Proteins in Membranes (OPM) database (*51*). The distance between the center of mass of bilayer and the center of mass of the protein was between 50 and 70 Å which corresponds to a minimum protein-bilayer distance between 5 and 25 Å for different simulations. Bilayers comprised 256 lipid molecules (128 lipids for each leaflet).

### Simulations protocol

All simulations were performed using NAMD (v 2.13) (*52*) with the CHARMM36 force field (*53*) and its CHARMM-WYF extension for the treatment of aromatics-choline interactions (*54*, *55*). After assembling the protein/membrane complex, the systems were first subjected to energy minimization with conjugate gradients (10000 steps). Then, six consecutive equilibrations were performed using the default equilibration protocol of CHARMM-GUI (*49*). During 50 ns equilibrations, we had gradual equilibrations of the initially assembled system; various restraints were applied to the protein, ligand, water, ions, and lipid molecules, similar to those used by Jo et al. (*56*). Next the production runs were performed for at least 2 μs using the coordinates and velocities of the last step of the equilibration run. All the production runs were done with an integration step of 2 fs in the NPT ensemble. Temperature and pressure were set at 310 K and 1 bar, respectively. Langevin dynamics with a temperature damping coefficient of 1.0 and the Langevin piston method with an oscillation period of 50 fs and a damping timescale of 25 fs were used to control the temperature and pressure, respectively. The ratio of the unit cell in the x-y plane was kept constant. The SHAKE algorithm was applied to constrain all bonds between hydrogen and heavy atoms, including those in water molecules to keep water molecules rigid (*57*). Electrostatic potentials were calculated using the Particle Mesh Ewald (PME) method (*58*). A Lennard-Jones switching function of 10-12 Å was used for van der Waals interactions. For each of the systems, we ran at least two series of simulations independently. The simulation conditions are summarized in Table S1. All simulations were uploaded to the Norwegian national infrastructure for research.

### Building the holo-Golgi-POPC system

We used the already formed complex in the holo-neutral simulation (Figure 6A) with a POPC tail in the cavity as a reference structure to guide the building of our model complex. First, we extracted a START structure with a ceramide molecule from our holo-Golgi simulation with already broken hydrogen bonds between the bound ceramide and residues at the binding site (Y482, N504, and Y553). We then aligned the model structure to the reference structure, transferred the POPC lipid from the reference holo-neutral simulation. Then a single point energy calculation has been performed using the CHARMM-GUI PDB Reader & Manipulator (*59*, *60*). This step ensured that the atomic coordinates were successfully defined and that the structure was ready to use in CHARMM-GUI Membrane Builder tool (*49*). The complex was placed onto the Golgi-like bilayer using the same orientation and penetration depth as observed in our other simulations. The system was then solvated in a water box and neutralized with 51 K^+^ ions. To ensure that the final system had no steric clashes or inappropriate geometry, an energy minimization was performed prior to starting a 2 μs-long dynamics simulation (and a replica) using the same protocol as above.

### Simulations trajectories analyses

The backbone RMSD between each simulation frame and the solvated and minimized X-ray structure was calculated using VMD (*61*) and is reported in supporting information (Figure S1). The RMSD was calculated for the bound form of proteins. The average RMSD for each of the six systems varies from 2.5 Å to 3.4 Å, a high value mostly accounted for by the mobile N-terminal helix in some simulations. The range of average RMSD values falls down to 2.1-2.6 Å if residues 362-392 are omitted from the calculation.

The electrostatics surface potential of START (apo) was calculated using APBS (*62*) and Pymol^30^. The protein tilt angle was defined as the angle between the bilayer normal and the long axis of the C-terminus α4 helix, defined by the alpha carbons of A565 and T591 (Figure S2). The angle was calculated using GROMACS analysis tools (*64*) for the whole 2 μs simulations. The electron density is calculated using VMD Membrane Plugin (*65*), and during the last 500 ns of each simulation. The depth of insertion of START domain is calculated using in-house Python code for each amino acid as the distance between its alpha carbon and the average plane of the phosphorus atoms. The average plane is calculated over the binding to the bilayer to end of the simulations.

Hydrophobic contacts between atoms are considered to exist if two unbound candidate atoms (Table S9) are within 3 Å for at least 10 ps. If one or more such contact is detected between atoms of two amino acids (or one aa and a lipid) we consider that these amino acids (or aa and lipid) engage in a hydrophobic contact (e.g. for plot on Figure 5). The criteria for hydrogen bonds are the following: acceptor (A) to hydrogen distance ≤2.4 Å and angle D−H−A (D: hydrogen-bond donor) ≥130°. These two criteria must be met for at least 100 ps. Cation−π interactions between the aromatic amino acids (W, F and Y) and lipids were considered to exist when all distances between the aromatic ring atoms and the choline nitrogen were below 7 Å. The candidate atoms for each category of interactions are listed in (Table S9). We used an in-house parallel Python3 code to perform the analysis. The code reads DCD trajectories files and can be run on modern multi-core computers. The code is based on MDAnalysis (*66*, *67*) for parsing of structure and trajectory files and for detection of hydrogen bonds (hydrogen bond analysis module). The most computing-intensive part, distance calculation for the detection of hydrophobic contacts, adopts a parallel implementation with Numba [https://numba.pydata.org/] and can be accelerated on multiple CPU cores. The code is available at (https://github.com/reuter-group/MD-contacts-analysis). The inventory of protein-lipid hydrophobic contacts, hydrogen bonds and cation-pi interactions were calculated for the bilayer-bound form of START, i.e. from the binding event and until 2 μs.

The gate opening is quantified using the distances between P564 (α4) and W473 (Ω4) in one hand, and W562 (Ω1) and S476 (Ω4) in the other hand. The lipid tail tilt angle with respect to Z axis were calculated using the vector defined between the C2 and C218/C316. The lipid tails snorkeling is evaluated based on the criterion that the last carbon in the tail is higher than 5 Å below the average phosphate plane (z > −5Å) for more than 1% of the 2 μs simulation.

### Free energy calculations

Calculations of relative binding free energy were performed using multisite lambda dynamics (MS*λ*D) (*68*, *69*). The calculations were performed with the 47a2 version of the CHARMM biomolecular package (*70*) using either DOMDEC (*71*) or BLaDE (*72*) on graphical processing units (GPUs) and the CHARMM36 force field. The protocols used for system preparation and simulation can be found in the supplementary information (Supplementary Text S2).

### Visualization

Visual analysis as well as image and movie generation were done using VMD (*61*) and PyMOL (*63*).

### Characterization of CERT-lipid complexes

CERT (isoform 2) was cloned in frame with an N-terminal His6-HA-StrepII-tag in a pcDNA5/FRT/TO vector for expression in HEK293 cells. HEK293 cells were maintained in DMEM, supplemented with 10% (v/v) FBS, 1% L-glutamine, and kept under selection in the presence of 15 µg/ml blasticidin and 100 µg/ml Zeocin before transfection. To create stable inducible human cell lines for LTPs, Flp-In T-Rex-293 cells were co-transfected with the plasmid coding for the tagged LTP and the pOG44 plasmid encoding the Flp recombinase (Invitrogen). Positive clones were selected by adding 100 µg/ml hygromicin B and 15 µg/ml blasticidin on the day after transfection. Cells were seeded and grown in the presence of 1µg/ml tetracycline (without other antibiotics) until 95% confluence, harvested, pelleted, and stored at −80°C for later use. CERT expression was evaluated by western blot with alpha-HA antibody, and all cell lines were tested for the presence of mycoplasma.

Cell lysis was performed by resuspending cell pellets in lysis buffer (50 mM Tris-HCl, 250 mM NaCl, 0.5 mM DTT; 2 µM avidin, protease inhibitors cocktail (Roche) and DNAse (Roche), at pH 7.4) and leaving on ice for 20 minutes. The final cell extract was obtained by centrifugation for 20 min at 16,000 g at 4°C in a bench top centrifuge, followed by centrifugation of the previous supernatant at 49,000 rpm, using a TLA100.4 rotor. Protein-lipid complexes were isolated via the Strep-II tag, at room temperature and eluted with 5 mM5 biotin. The eluted complexes were centrifuged at 16,000 g for 5 minutes and fractionated on a Superdex 200 SEC column (Invitrogen). Proteins and lipids in the SEC fractions were analyzed by sodium dodecyl sulfate polyacrylamide gel electrophoresis (SDS-PAGE) on pre-cast 4-12% gradient gel (Life technologies) and by LC-MS/MS, respectively.

Lipids were separated on an Agilent 1260 HPLC system consisting of a degasser, binary pump, and autosampler directly coupled to a Q-Exactive Plus (Thermo) equipped with a heated ESI source. The column was a Kinetex 30×2.1 mm, 2.6 µm, C18 100 Å (Phenomenex). A binary solvent system was used in order to separate lipids. The mobile phase A consisted of H_2_O:acetonitrile (60:40), 10 mM ammonium formate, and 0.1% formic acid, while the mobile phase B consisted of isopropanol:ACN acetonitrile (90:10), 10 mM ammonium formate, and 0.1% formic acid. The separation started at 80% buffer A and 20% buffer B. In a 3 min gradient, buffer B was increased from 20% to 50%, followed by a 10 min gradient from 50% to 70% buffer B. Finally, a 5.4 min gradient was applied to increase buffer B from 70% to 97%. The column was subsequently washed for 2.1 min with 97% buffer B and equilibrated for 3.6 min with 20% buffer B. The flow rate was 500 µl/min. The entire run was 24 min long. The effluent was directly introduced into the ESI source of the MS, and analyzed each time for each paired fraction in either positive or negative ionization mode. The ESI source ion spray voltage was set to 1.7 kV. The mass spectrometer was operated in the mass range from 250 to 1,600 m/z. Charge state screening was not enabled. The ten most abundant peaks (TOP 10) were selected and fragmented in MS2 by HCD. The normalized collision energy (NCE) was 30.

## Acknowledgments

We are grateful to members and former members of ACG’s group at the Department of Cell Physiology and Metabolism, University of Geneva for continuous discussions and support.

## Author contributions

Conceptualization: M.M., N.R., and A.C.G. Methodology: M.M., N.R., and A.C.G. Acquisition and analysis of data: M.M., P.G., R.T, A.C., K.T. Visualization: M.M Supervision: M.M., N.R., and A.C.G. Writing—original draft: M.M. Writing—review and editing: M.M., K.T., N.R., and A.C.G. All authors discussed the results and participated in interpreting the results.

## Data and materials availability

All the MD trajectories are uploaded to the Norwegian national infrastructure for research data (NIRD), have been issued a DOI (10.11582/2023.00139) and can be accessed using the following URL: https://doi.org/10.11582/2023.00139

## Supplementary Materials

**This PDF file includes:**

Supplementary Text

Figs. S1 to S15

Tables S1 to S9

**Other Supplementary Materials for this manuscript include the following:**

Movies S1 to S4

