## Supporting Information for "A Mechanistic Model for the Release of Ceramide from the CERT START Domain"

Mahmoud Moqadam *et al.*

**This PDF file includes:**

Supplementary Text
Figs. S1 to S15
Tables S1 to S9

**Other Supplementary Materials for this manuscript include the following:**
Movies S1 to S4

### Supplementary Text.

#### S1. Replicate MD Simulations

We conducted two replicates of the simulations for each START-lipid bilayers systems (see Table S1); the results of the first replicates are presented in the main text. In this section, we report our observations from the second replicates. Our findings suggest that only POPC can insert a tail into the cavity, positioning its head group between the Ω1 and Ω4 loops.

***Apo-ER_r2.*** A POPE lipid occupied the site between Ω1 and Ω4, and the cavity remained closed throughout the simulation. No gate opening was observed during the simulation.

***Apo-neutral_r1.*** a POPC lipid occupied the site between Ω1 and Ω4 snorkeling to insert a tail in the open cavity. Simultaneously another POPC lipid blocked the entrance into the cavity by placing its headgroup near the cavity. No tail insertion was observed during the simulation.

***Holo_neutral_r2 / holo-Golgi_r2.*** we also observed the opening of the cavity and lipid snorkeling events. In both simulations, a POPC occupied the site between Ω1 and Ω4 loops and snorkels around the entrance to the cavity, but neither tail insertion nor ceramide release were observed.

#### S2. Free energy calculations, simulation protocols

1. **System preparation.**

***START apo and START-Cer.*** The structure of the START-Cer complex was retrieved from the protein data bank (PDB ID: 2e3q) and processed with MMTSB (*73*) tools and CHARMM-GUI. The CHARMM simulation package (*74*) (v47a2) was then used with the CHARMM36 force field for proteins and lipids to generate the topology (psf) and coordinates (crd) files for the complex. The complex was energy-minimized (steepest descents: 100 steps, adopted basis Newton-Raphson: 1000 steps), solvated in a cubic box (a=b=c= 86.1 Å) with TIP3P water molecules (edge cutoff=10 Å) and neutralized by adding 3 counter $K^+$ ions, replacing 3 randomly chosen water molecules from the bulk. This was followed by energy-minimization (steepest descent: 200 steps, adopted basis Newton-Raphson: 1000 steps). Particle mesh Ewald (PME) was used for long-range electrostatic interactions and periodic boundary conditions with a non-bonded cutoff of 16 Å along with truncation (vswitch) of van der Waals interactions at 10 Å. The system was then gradually heated from 190 K to 310 K with increments of 1 K every 100 steps using a gaussian distribution for velocity assignments. SHAKE (*75*) was used to constrain bonds between heavy atoms and hydrogen atoms. A short 5 ns equilibration and then 400 ns production simulations were run in the NPT ensemble with a 2-fs integration time step and using BLaDE (*72*) in CHARMM 47a2. The temperature was set to 310 K and controlled using a Langevin thermostat (drag coefficient = 0.1 $ps^{-1}$). A Monte Carlo barostat was used (target pressure=1 atm) with pressure coupling moves every 25 steps and the default (100 $Å^3$) standard deviation of the gaussian distribution was used for drawing volume changes.

The trajectory obtained was recentered around the START domain using CHARMM v.47a2 and clustered, using ttclust (*76*) and the RMSD of the domain backbone atoms. A structure from the largest cluster was selected for free energy calculations. Cer was removed from that structure to generate the structure of apo START (except for the N504 mutations for which the cluster closer to the X-ray structure was used).

***START-Cer-POPC and START-POPC.*** The structure of the START-Cer-POPC complex was taken from the holo-Neu_r1 simulation. The conformation was chosen such that the Cer and POPC tails were present and before Cer leaves the binding site (t=886.9 ns). The START-POPC structure was obtained from the above by removing Cer.

#### 2. Multisite lambda dynamics (MS$\lambda$D)

The apo START structure and the START-Cer complex structure extracted from the cluster above were solvated (cutoff 15 Å) in a cubic box with TIP3 water molecules and neutralized by adding three K+ ions using the MMTSB toolset. These systems are denoted as **START-Water** and **START-Cer-Water** respectively in Table **S5**. The structures extracted from the holo-Neu r1 simulation were also similarly solvated and neutralized and are referred as **START-Cer-POPC-Water** and **START-POPC-water** in Table **S6**.

The mutant atoms on the perturbation site were added using the PATCH command in the CHARMM program (*68*). The perturbation site had all the atoms of the wild-type residue and all the atoms for the mutant (including backbone) (*69*). All valence angles, dihedrals and impropers were removed between the alchemical groups. The alchemical system containing two substituents on a single site was set up using the BLOCK module (*7*) of CHARMM and atoms of both substituents were constrained using the SCAT (force constant = 118.4) facility. A FNEX value of 5.5 was used for the functional form of $\lambda$. Soft core potentials were used along with the ALF (*77*) biases for each of the substituents. The POPC tail outside the START cavity was harmonically restrained with a force constant of 2 kcal*mol$^{-1}$*Å$^{-2}$ on heavy atoms.

The START-Water and START-Cer-Water systems were simulated using BLaDE with $\lambda$ values saved every 20 fs and a non-bonded cutoff of 12 Å along with truncation (vswitch) of van der Waals interactions between 9-10 Å while keeping the remaining conditions and parameters same as for the equilibrium simulations described earlier in this section (Cf *System preparation)*. The START-Cer-POPC-Water and START-POPC-Water systems were simulated with DOMDEC (*71*) since BLaDE does not support harmonic restraints. In these simulations the Nosé-Hoover thermostat (temp = 310 K; tmass = 1000 kcal*ps$^2$) was used for temperature control and a Langevin piston (friction coefficient: 20 ps$^{-1}$) for pressure control. All the simulations either with BLaDE or DOMDEC were performed with CHARMM 47a2.

MS$\lambda$D simulations were performed in an iterative fashion (*78*, *79*). The first phase consisted of 50-60 small 100 ps long simulations to get an idea regarding the biases and, the second phase consisted of 10-20 longer 1 ns simulation to flatten the biases. The ALF biases were calculated using weighted histogram analysis method (WHAM) (*80*) and optimized after every run. The production runs were started with the lambda landscape flattened in 5 independent replicas of 5 ns each. The final relative free energy differences are obtained via the following equation:

$$\Delta\Delta G = -k_B T ln(\frac{P(\lambda_j = 1)}{P(\lambda_i = 1}) - (\phi_j - \phi_i) \quad (1)$$

Where, i and j are the wild-type and mutant residues respectively.

P – probability

$\phi$ – bias free energy

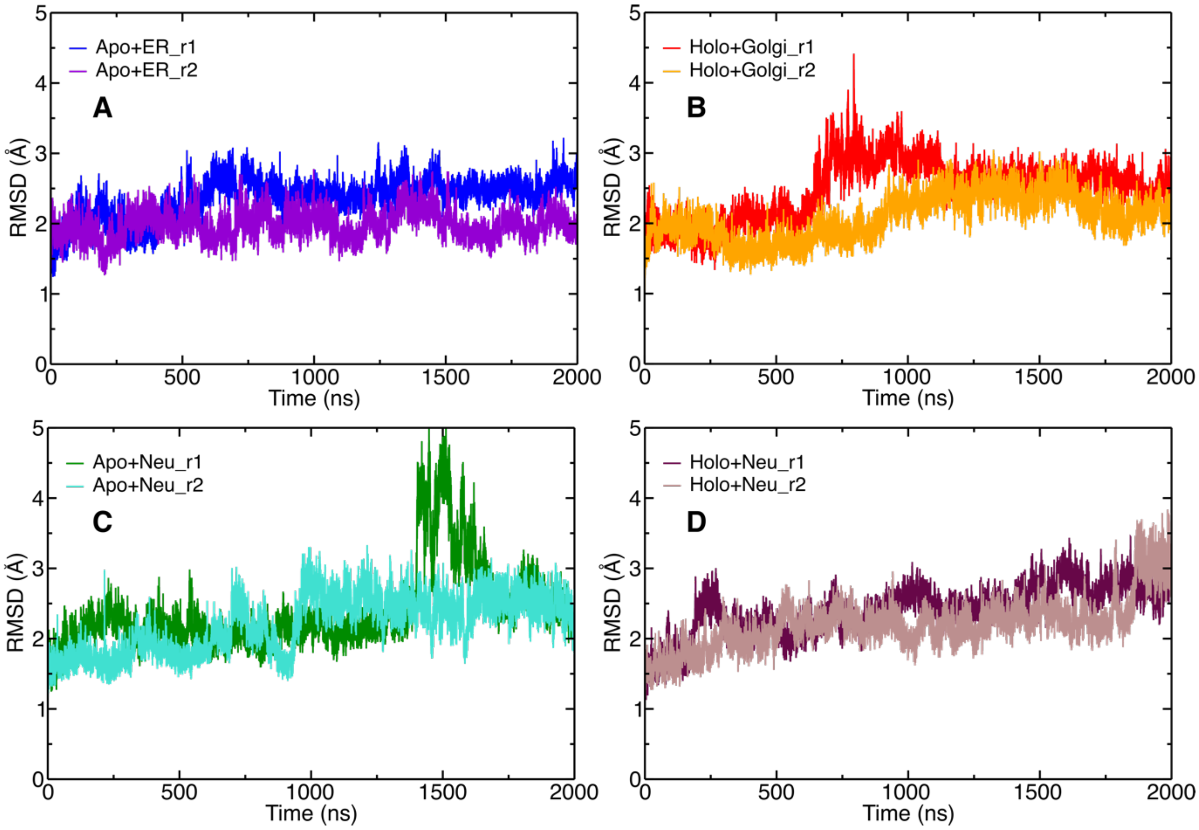


**Fig. S1. RMSD plots.** RMSD for both replicas (r1 and r2) in each case, calculated for START domain excluding the N terminus helix, resid 362-391. (**A**) Apo in the ER-like bilayer (**B**) Holo in the Golgi-like bilayer (**C**) Apo in the neutral bilayer (**D**) Holo in the neutral bilayer.

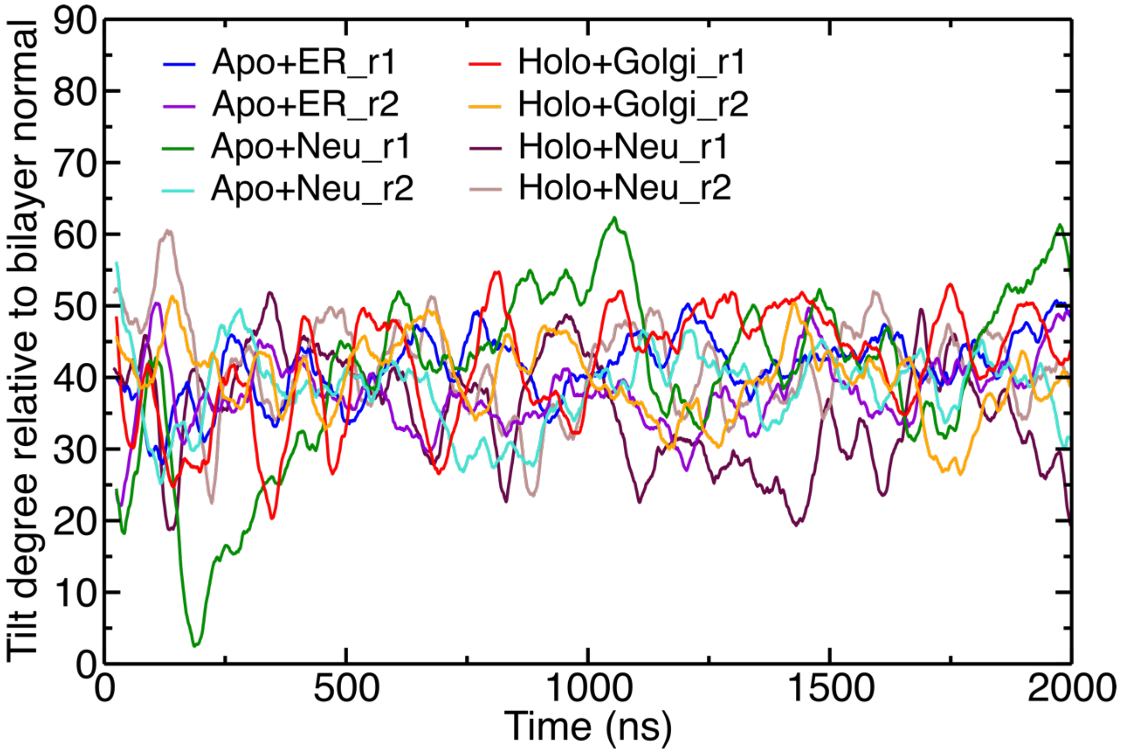


**Fig. S2. Tilt angle of apo and holo forms on the different bilayers.** The tilt is defined using the START domain α4 vector with respect to the bilayer normal.

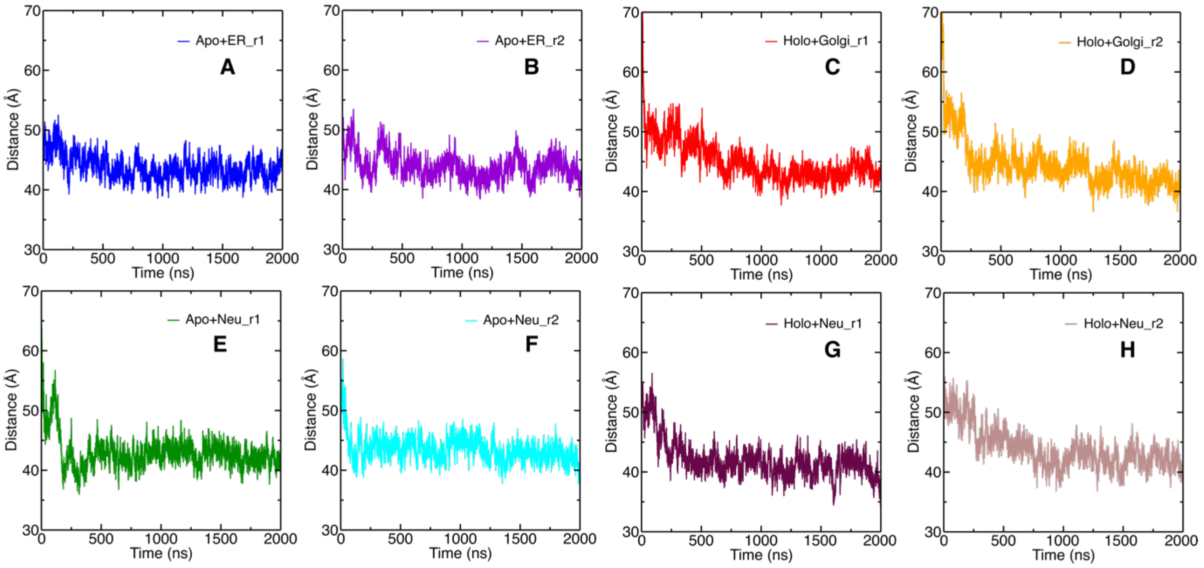


**Fig. S3. START domain and bilayer distance.** Distance between center of mass of the START domain and center of mass of the bilayer through the 2 $\mu$s simulations. (**A and B**) apo and ER bilayer. (**C and D**) holo and Golgi bilayer. (**E and F**) apo and neutral bilayer. (**G and H**) holo and neutral.

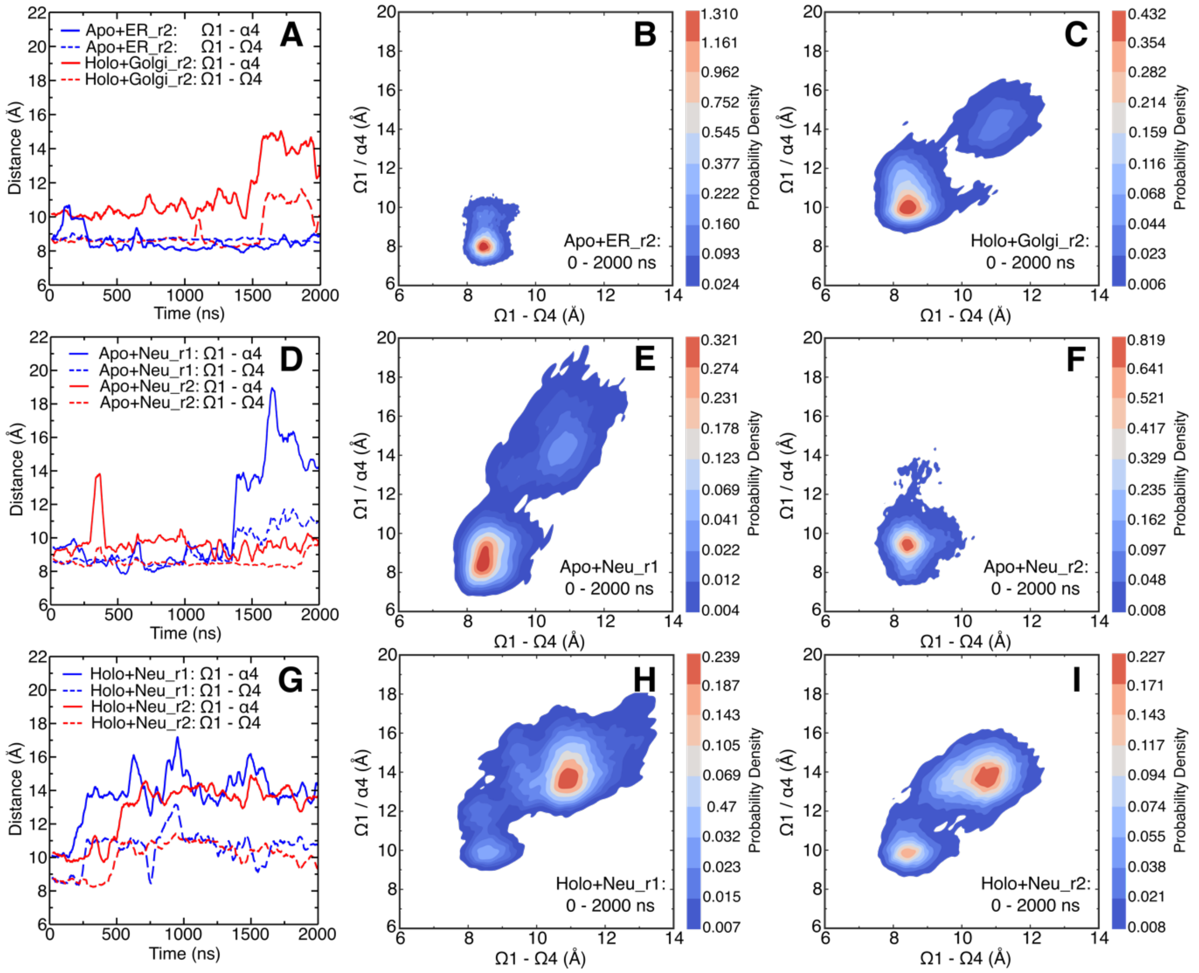


**Fig. S4. Gate opening through displacements of Ω1, Ω4 and α4.** (**A, D, and G**) Time series of distances between Ω1and α4 (plain lines), and Ω1 and Ω4 (dotted lines) in simulations of START with the ER (**A**), Golgi (**A**), and neutral (**D, G**) bilayers. (**B, C, E, F, H, and I**) Density distributions of the closed and open states of START simulated with the ER bilayer (**B**), the Golgi bilayer (**C**), the neutral bilayer (Neu) (**E, F, H, and I**).

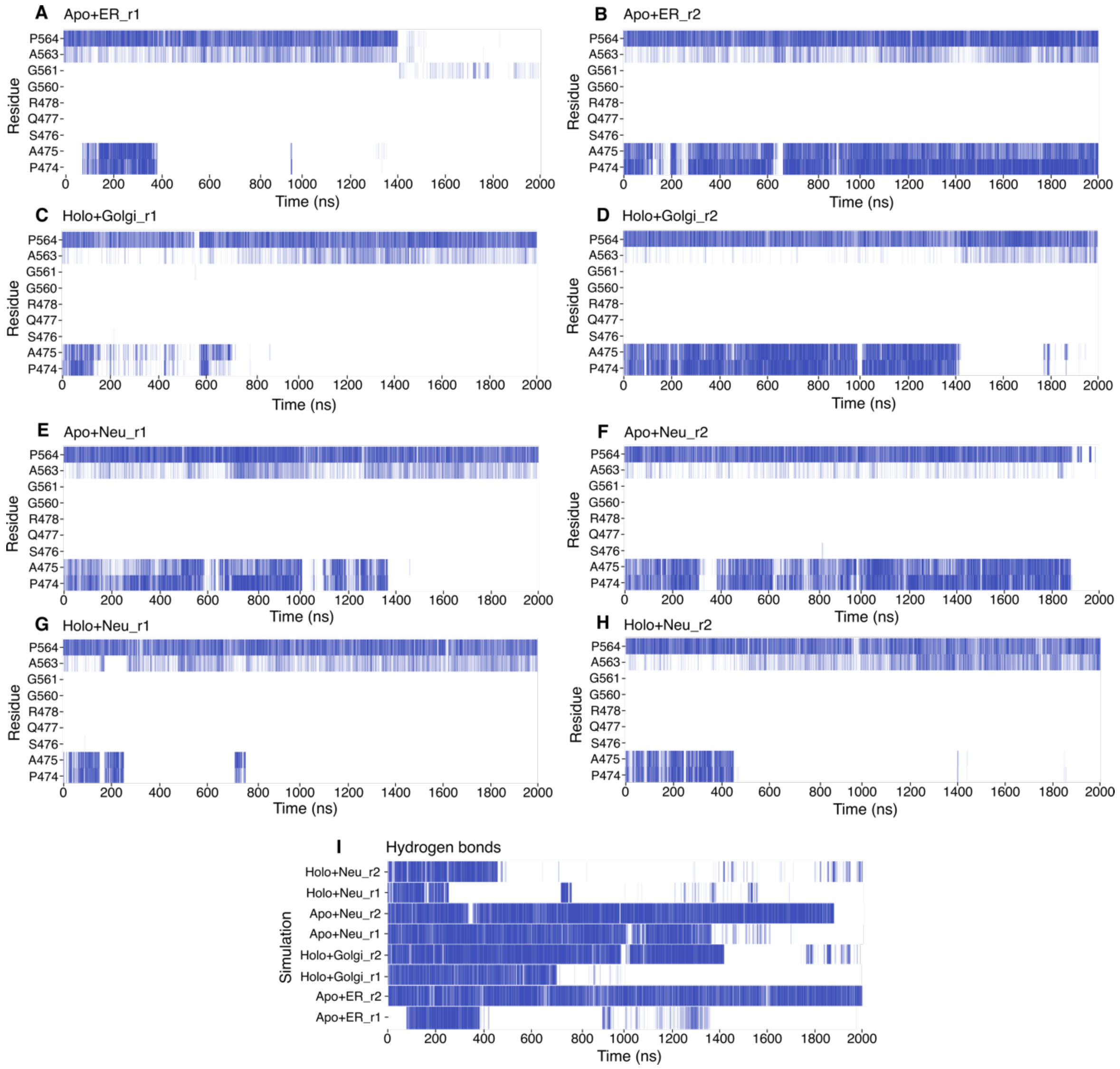


**Fig. S5. Hydrophobic contacts and hydrogen bonds.** Time series of hydrogen bonds and hydrophobic contacts between W562 and its nearest neighbors in the Ω1 and Ω4 loops. (**A-H**) Hydrophobic contacts (**I**) Hydrogen bonds. The plots display a bar if there is at least one hydrophobic contact (A-H) or hydrogen bond (I).

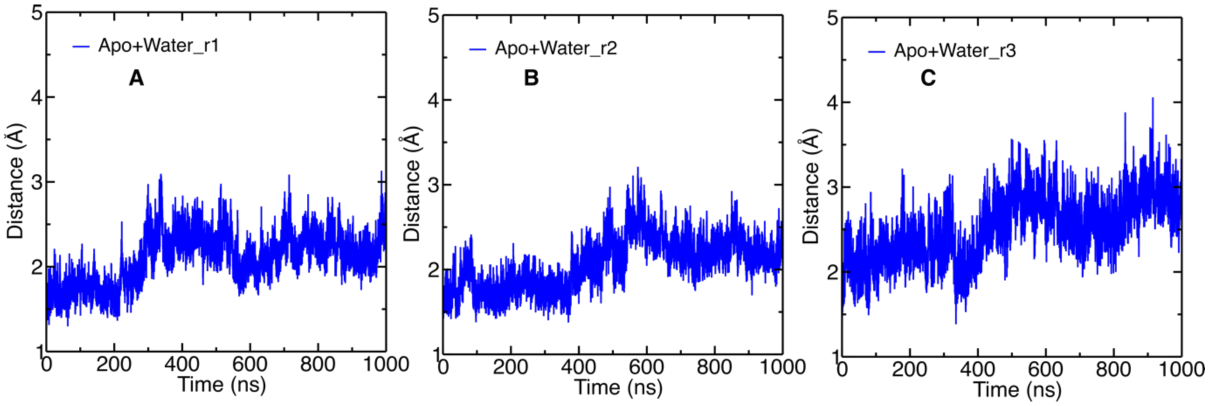


**Fig. S6. Backbone RMSD for apo START in water in three replicate simulations.** The N-terminus helix (residues number 362-391) is not included.

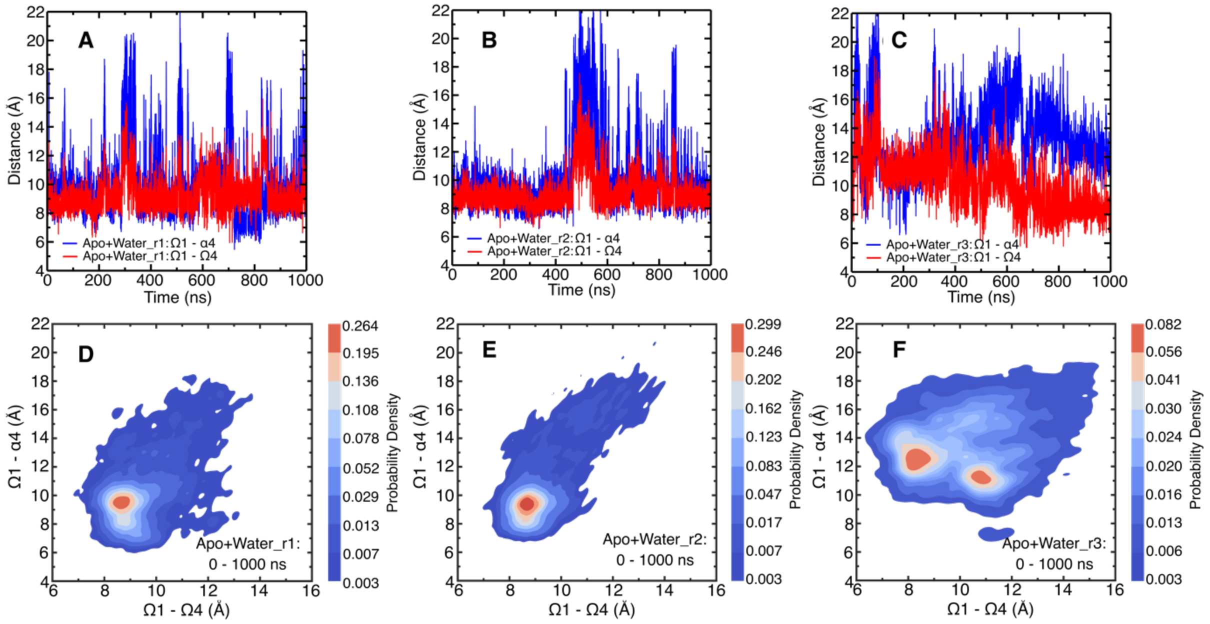


**Fig. S7. Gate opening through displacements of Ω1, Ω4 and α4.** (**A-C**) Time series of the Ω1-α4 (blue) and Ω1-Ω4 (red) distances in each of the three replicate simulations of apo START in water. (**D-F**) Density distributions of the distances plotted in A-C and characterizing the closed and open states of START.

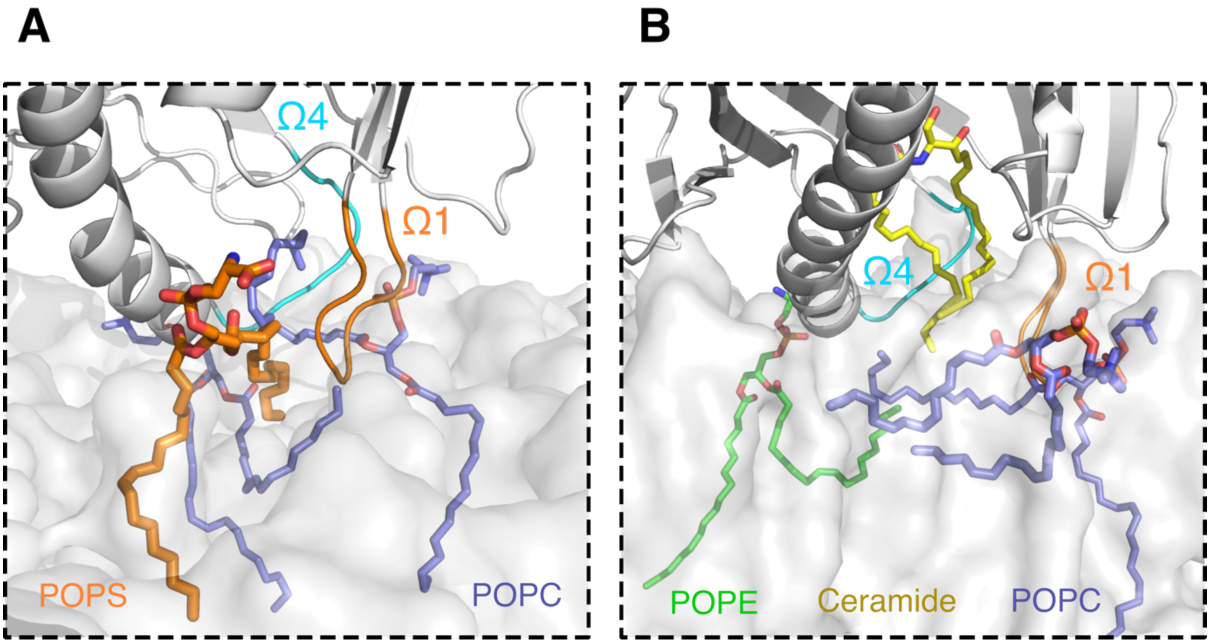

**Fig. S8. Lipid Snorkeling.** Snapshots illustrating lipid snorkeling under the START domain in (**A**) the ER-like and (**B**) the Golgi-like bilayers.

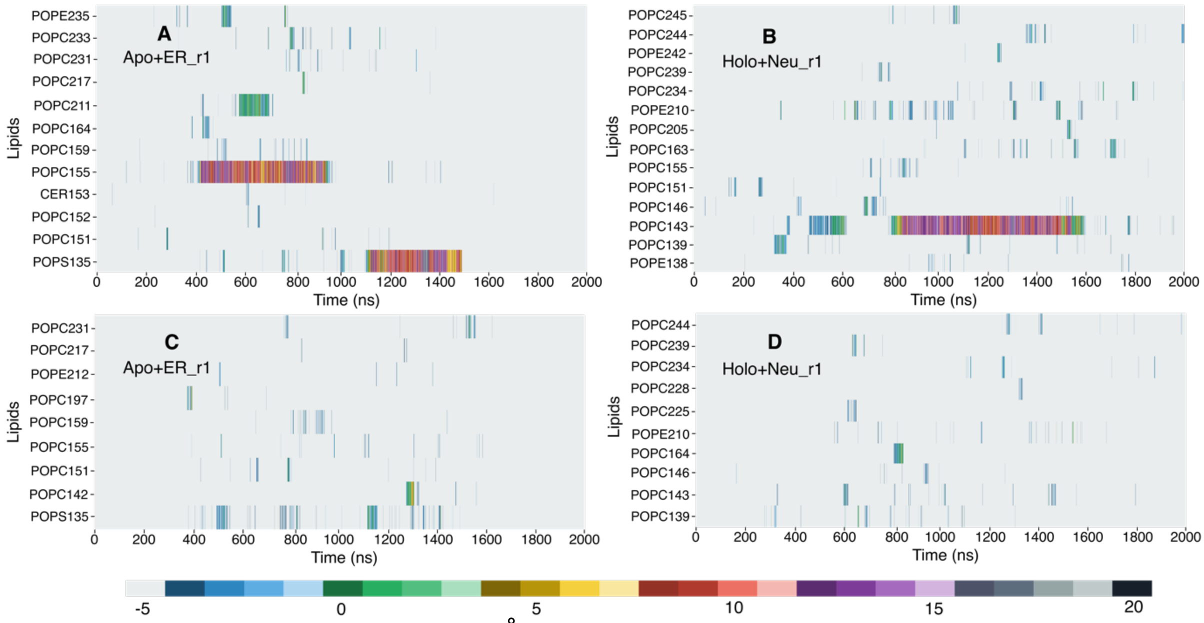


**Fig. S9. Lipid snorkeling within 3Å of the START domain in the ER and neutral simulations.** Z value of the C218 of the POPX (X: C, E, I, and S) and the C18S atoms of the ceramide with respect to the average plane of the phosphorus atoms (Z=0) (**A**) apo+ER_r1 (**B**) holo+Neu_r1. Z value of the C316 of the POPX (X: C, E, I, and S) and the C18F atoms of the ceramide with respect to the average phosphate plane (Z=0) (**C**) apo+ER_r1 (**D**) holo+Neu_r1.

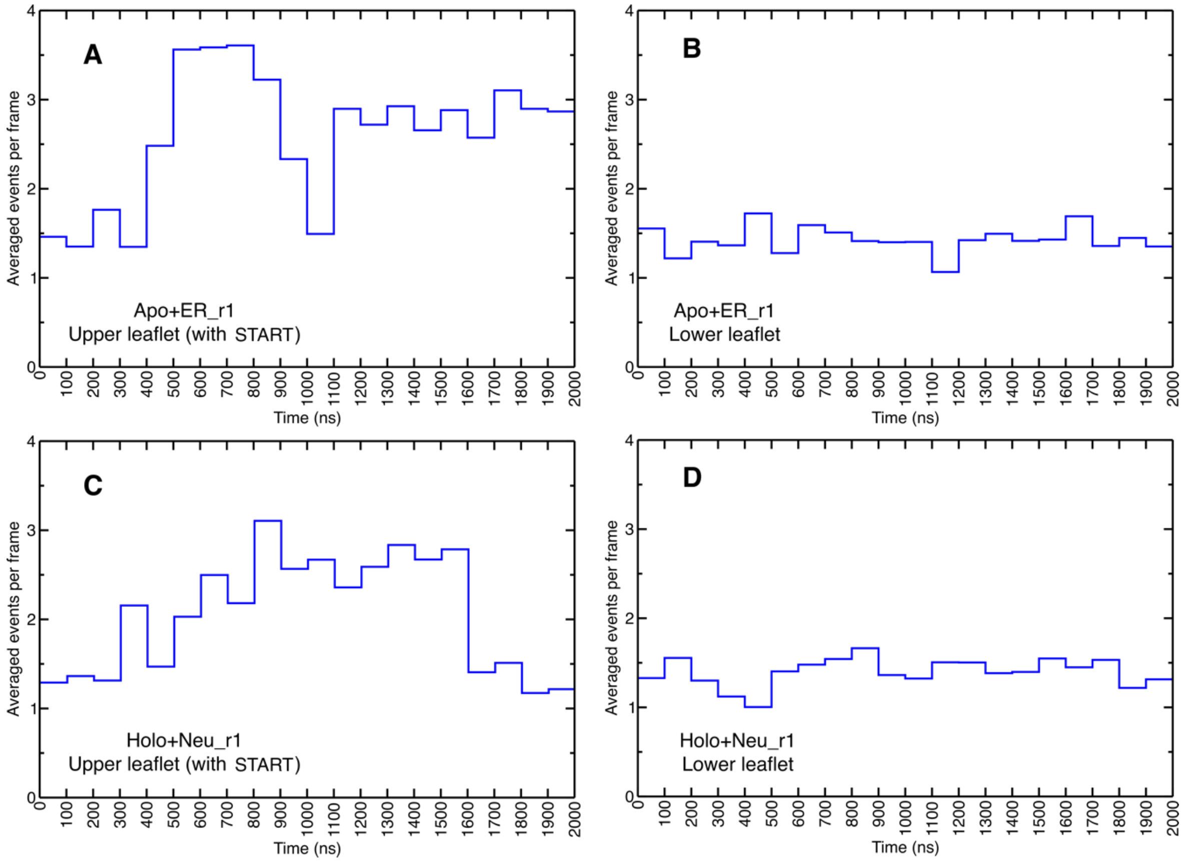


**Fig. S10. Average number of snorkeling events.** Number of time (average per frame) the C218 and C316 atoms of the POPX (X: C, E, I, and S) and the C18S and C18F atoms of the ceramide are at least 5 Å higher than the average phosphate plane. Data shown for lipids from (**A**) the upper leaflet (to which the START domain is bound) and (**B**) the lower leaflet in the apo+ER_r1 simulation; and for lipids in (**C**) the upper leaflet (with the START domain) and (**D**) the lower leaflet in the holo+Neu_r1 simulation. The lower leaflets are shown for reference.

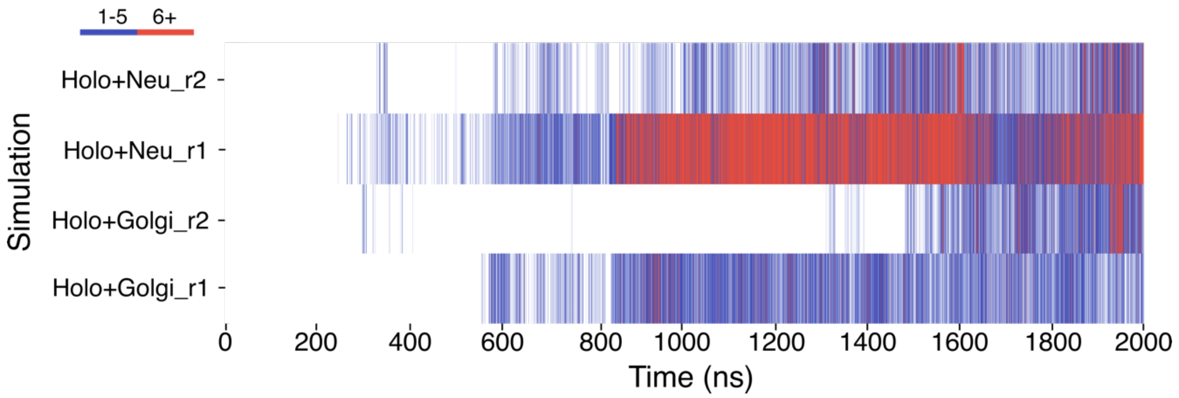


**Fig. S11. Hydrophobic contacts between bound ceramide and lipid bilayer.** Number of hydrophobic contacts between the START-bound ceramide and the lipids of the bilayer in the holo+Golgi and Holo+Neu simulations. Color bar indicates the number of hydrophobic contacts in each frame; Blue: 1-5 hydrophobic contacts, red: 6 or more hydrophobic contacts.

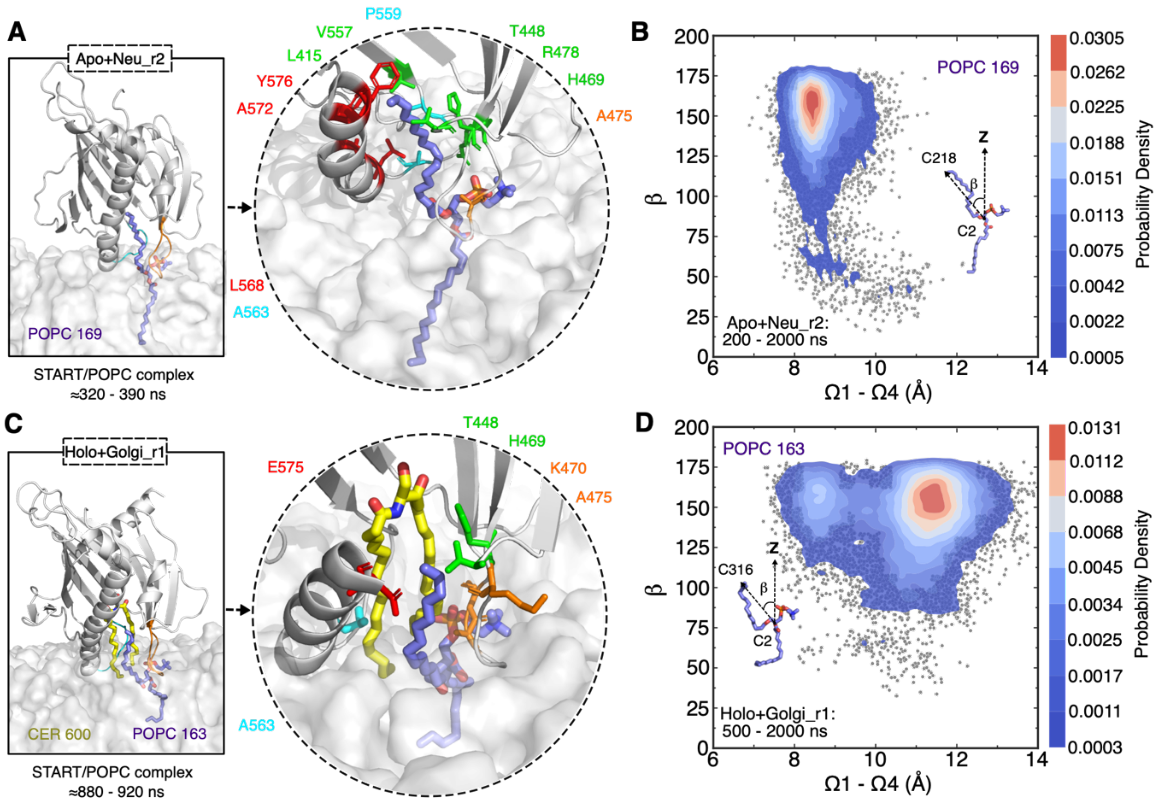


**Fig S12. POPC (1-palmitoyl-2-oleoyl-phosphatidylcholine) rearrangement induced by START domain in opened state detected by MD simulation.** Close-up view of the binding conformation of POPC within apo START domain, and the amino acids in the cavity involved in hydrophobic contacts with the POPC tail in the neutral bilayer. (**B**) Density distribution of the POPC tail angle with respect to membrane normal in the neutral bilayer is presented. (**C**) Close-up view of the binding conformation of POPC within START-Cer complex, and the amino acids in the cavity involved in hydrophobic contacts with the POPC tail in the Golgi bilayer. (**D**) Density distribution of the POPC tail angle with respect to membrane normal in the Golgi bilayer is presented. The times at which the POPC tail inserts into and exits the cavity are given below the snapshots. Ceramide colored yellow, POPC colored purple, Ω1 colored orange and Ω4 colored cyan. START domain is shown as cartoon and colored grey. Membrane is shown as grey transparent surface. The gray dots represent all the sampled tilt angles.

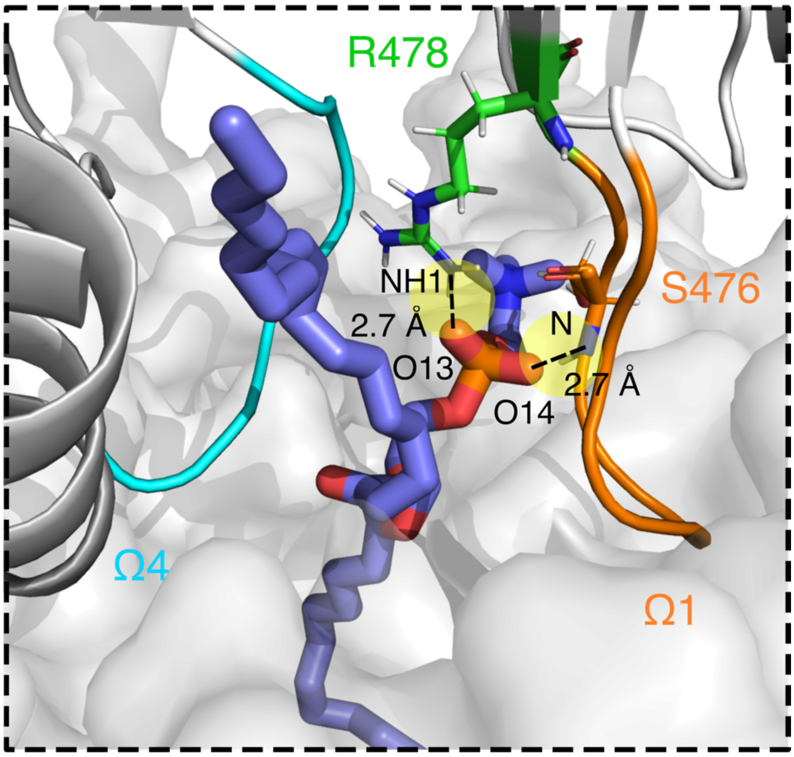


**Fig. S13. The POPC lipid located between Ω1 and Ω4.** The POPC head group is locked in place by a salt bridge between its phosphate group and R478, and by hydrogen bonds between the S476 side chain and the POPC phosphate group.

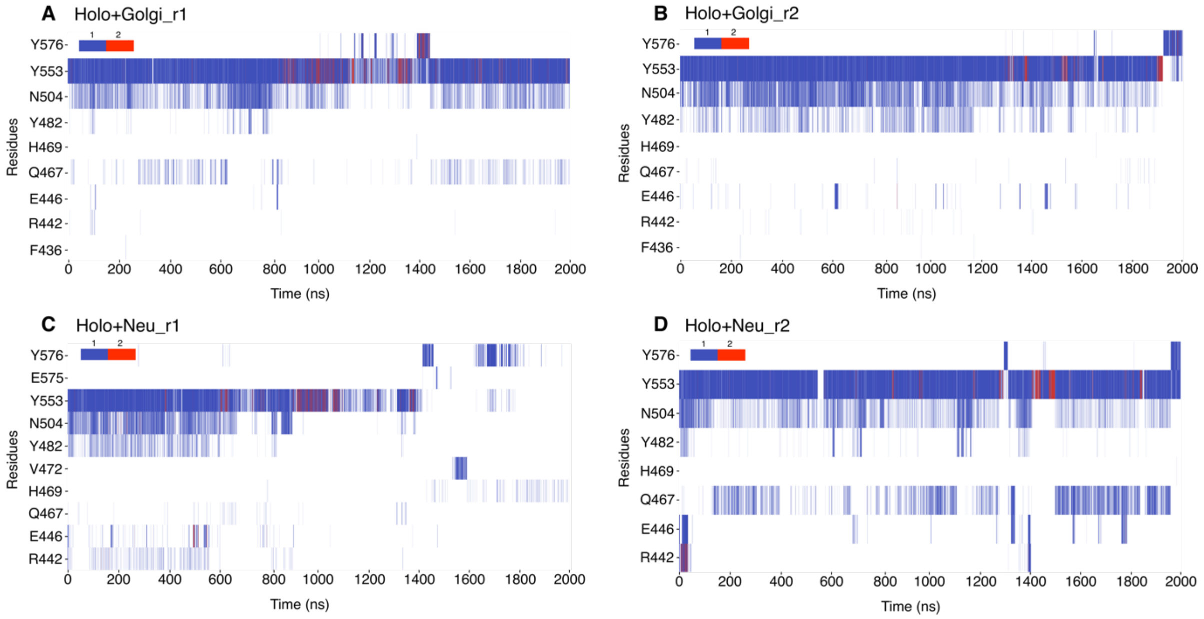


**Fig. S14. Hydrogen bonds network around the ceramide head group.** Time series along the Holo+Golgi_r1 (**A**), Holo+Golgi_r2 (**B**), Holo+Neu_r1 (**C**) and Holo+Neu_r2 (**D**) simulations. Color bar indicates the number of hydrogen bonds in each frame; Red: 2 hydrogen bonds, blue: 1 hydrogen bond.

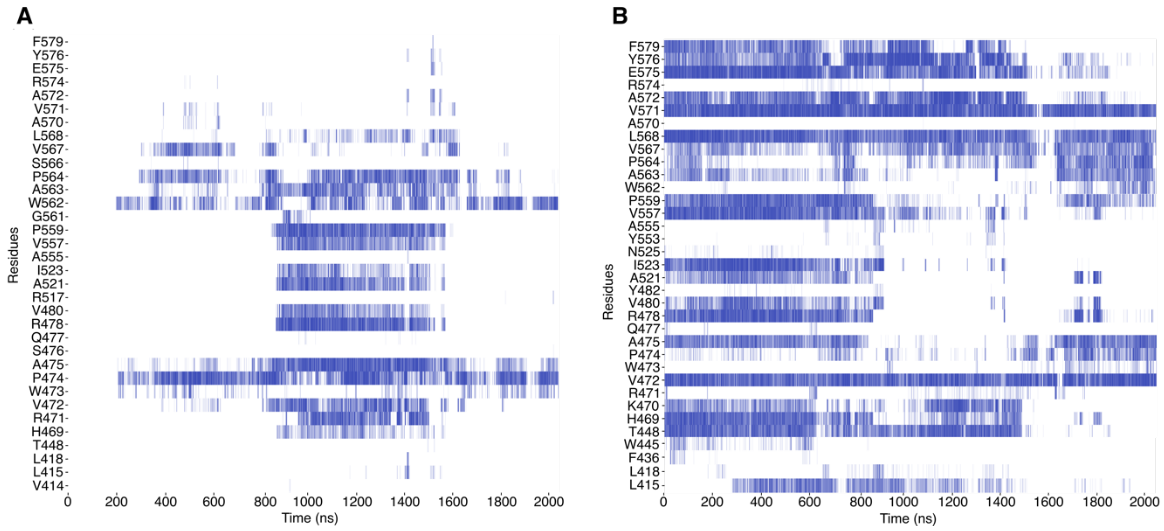


**Fig. S15. All hydrophobic contacts in the holo+Neu_r1 simulation.** Time series of contacts between (**A**) POPC143 and the START domain, between (**B**) ceramide and the START domain.

| # | Name | Domain | Lipid bilayer | Total number of atoms | Number of water molecules | Ions | Initial cell dimensions | SA/Lipid ($Å^2$) | Binding at (ns) |
|---|---|---|---|---|---|---|---|---|---|
| 1 | Apo+Water_r1 | Apo | - | 66858 | 21023 | 3 $K^+$ | 89.0<br>89.0<br>89.0 | - | - |
| 2 | Apo+Water_r2 | | | | | | | | |
| 3 | Apo+Water_r3 | | | | | | | | |
| 4 | Apo+ER_r1 | Apo | 53% POPC<br>23% POPE<br>10% POPI<br>5% POPS<br>5% CHL<br>4% Cer | 135953 | 33110 | 43 $K^+$ | 90.3<br>90.3<br>176.3 | 59.6 | 300 |
| 5 | Apo+ER_r2 | | | 168992 | 44123 | | 90.3<br>90.3<br>218.3 | 59.6 | 500 |
| 6 | Apo+Neu_r1 | Apo | 60% POPC<br>30% POPE<br>5% CHL<br>4% Cer | 147012 | 36931 | 3 $K^+$ | 89.9<br>89.9<br>191.3 | 58.4 | 400 |
| 7 | Apo+Neu_r2 | | | 169467 | 44416 | | 89.9<br>89.9<br>220.3 | 58.4 | 200 |
| 8 | Holo+Neu_r3 | Holo | 60% POPC<br>30% POPE<br>5% CHL<br>4% Cer | 143500 | 35731 | | 89.9<br>89.9<br>186.9 | 58.5 | 200 |
| 9 | Holo+Neu_r4 | | | 143500 | 35731 | | 89.9<br>89.9<br>161.9 | 58.5 | 300 |
| 10 | Holo+Golgi_r1 | Holo | 46% POPC<br>18% POPE<br>9% POPI<br>6% POPS<br>5% POPA<br>5% DAG<br>5% CHL<br>6% Cer | 123852 | 29203 | 51 $K^+$ | 89.9<br>89.9<br>161.9 | 58.8 | 500 |
| 11 | Holo+Golgi_r2 | | | 123852 | 29203 | | 89.9<br>89.9<br>186.9 | 58.8 | 300 |
| 12 | Holo+Golgi +POPC_r1 | Holo + POPC | | 142813 | 35471 | | 91.0<br>91.0<br>176.3 | 59.0 | Already bound structure |
| 13 | Holo+Golgi +POPC_r2 | | | 142813 | 35471 | | 91.0<br>91.0<br>176.3 | 59.0 | Already bound structure |

**Table S1. Composition and size of the simulated systems.** Each was simulated twice for 2 µs (production run). The raw trajectories are available at the NIRD Research Data archive (DOI: 10.11582/2023.00139).

| SSE | AA | APO | | HOLO | |
|---|---|---|---|---|---|
| | | ER | Neu | Golgi | Neu |
| **Hydrophobic contacts (avg. per frame)** | | | | | |
| Ω1 | V472 | 1.8 | 1.0 | 1.0 | 1.6 |
| | W473 | 5.5 | 5.4 | 4.9 | 5.2 |
| | P474 | 4.0 | 3.4 | 3.0 | 4.0 |
| Ω4 | W562 | 2.0 | 2.2 | 2.4 | 2.8 |
| | P564 | 1.3 | 1.0 | 1.3 | 1.9 |
| α4 | S566 | 1.1 | 1.3 | 1.0 | 1.1 |
| | V567 | 3.6 | 3.0 | 2.2 | 3.0 |
| | V571 | 1.8 | 1.5 | 1.4 | 1.6 |
| | R574 | 0.6 | 0.7 | 0.6 | 0.7 |
| **Hydrogen bonds (%)** | | | | | |
| Ω1 | R471 | 67.0 | 49.2 | 68.5 | 68.8 |
| | W473 | 48.4 | 41.0 | 56.3 | 50.4 |
| | S476 | 64.9 | 43.3 | 60.8 | 64.9 |
| β6 | Q477 | 30.1 | - | - | - |
| | R478 | 60.1 | 30.2 | 34.1 | 56.1 |
| Ω2 | R517 | 61.0 | 37.0 | 66.4 | 56.4 |
| Ω4 | W562 | 53.2 | 41.4 | 46.5 | 38.0 |
| α4 | A565 | 36.8 | 32.1 | 45.2 | 44.1 |
| | S566 | 62.3 | 48.8 | 37.3 | 51.6 |
| | R569 | - | 42.5 | 41.3 | 42.2 |
| | K573 | 31.5 | 30.0 | 38.5 | 31.0 |
| | R574 | 63.2 | 50.9 | 67.8 | 65.7 |
| | K578 | 45.6 | - | 44.6 | 36.5 |
| | R582 | 38.6 | - | - | - |
| **Cation-π interactions (%)** | | | | | |
| Ω1 | W473 | 11.3 | 5.5 | 1.1 | 3.6 |
| Ω4 | W562 | 19.5 | 17.7 | 20.2 | 13.4 |

**Table S2: Inventory of interactions between START and the bilayer lipids.** The data presented in the table are averages over two replicas in each case, calculated from binding to the bilayers to the end of the simulations. SSE: Secondary structure elements. AA: amino acid, α: helix, Ω: loop, β: beta-strand, occ: occupancy. Average number of hydrophobic contacts per frame are reported if > 0.5. Hydrogen bond occupancies are reported if > 30%. All cation-pi interactions are reported.

| Simulations | W473 | | | W562 | | |
|---|---|---|---|---|---|---|
| | Phosphate | Glycerol | Choline | Phosphate | Glycerol | Choline |
| Apo+ER_r1 | 22.9 | 71.9 | 5.2 | 87.7 | 8.2 | 4.1 |
| Apo+ER_r2 | 39.1 | 48.0 | 12.9 | 90.5 | 7.3 | 2.2 |
| Holo+Golgi_r1 | 33.7 | 62.7 | 3.6 | 88.0 | 11.3 | 0.7 |
| Holo+Golgi_r2 | 32.1 | 56.7 | 1.0 | 65.1 | 22.5 | 3.2 |
| Apo+Neu_r1 | 34.0 | 65.5 | 0.5 | 69.6 | 13.3 | 17.1 |
| Apo+Neu_r2 | 14.3 | 85.6 | 0.1 | 83.8 | 16.1 | 0.1 |
| Holo+Neu_r1 | 15.0 | 84.5 | 0.5 | 16.1 | 83.6 | 0.3 |
| Holo+Neu_r2 | 42.9 | 54.8 | 2.3 | 93.5 | 6.4 | 0.1 |

**Table S3. Occupancies (%) of lipid-tryptophane (W473, W562) hydrogen bonds.** Hydrogen bonds between the two exposed tryptophanes and the lipid bilayers, reported in percentage of simulation time.

| Residues | Holo+Golgi_r1 | Holo+Golgi_r2 | Holo+Neu_r1 | Holo+Neu_r2 |
|---|---|---|---|---|
| PHE-436 | 0.02 | 0.04 | 0.00 | 0.00 |
| ARG-442 | 0.18 | 0.19 | 0.15 | 2.33 |
| GLU-446 | 0.39 | 2.45 | 2.46 | 4.29 |
| GLN-467 | 8.41 | 0.31 | 0.37 | 28.08 |
| HSD-469 | 0.02 | 0.01 | 0.74 | 0.01 |
| VAL-472 | 0.00 | 0.00 | 1.77 | 0.00 |
| TYR-482 | 1.56 | 16.16 | 6.02 | 3.58 |
| ASN-504 | 31.71 | 47.06 | 20.97 | 28.43 |
| TYR-553 | 93.11 | 94.12 | 58.36 | 95.14 |
| GLU-575 | 0.00 | 0.00 | 0.24 | 0.00 |
| TYR-576 | 3.72 | 3.89 | 7.99 | 2.69 |

**Table S4. Occupancies (%) of Cer-START hydrogen bonds**. Inventory of hydrogen bonds between ceramide and residues of the START domain cavity in simulations of the holo START domain on Golgi-like and neutral bilayers. Numbers are occupancies reported in percentage of simulation time.

| Mutation | ΔG (kcal/mol) | | $\Delta\Delta G_{binding}$ (kcal/mol) | Production run |
|---|---|---|---|---|
| | START-Cer-Water | START-Water | START-Cer | (ns) |
| Y553F | 17.7 ± 0.1 | 16.9 ± 0.1 | 0.8 ± 0.1 | 200 |
| N504A | 81.5 ± 0.2 | 80.5 ± 0.2 | 1.1 ± 0.2 | 200 |
| V480A | 0.7 ± 0.1 | -0.5 ± 0.1 | 1.2 ± 0.1 | 200 |

**Table S5. MSλD START-Cer relative binding free energy (ΔΔG) for the Y553F, N504A and V480A substitutions.** The production simulations were run in five replicates (40 ns each).

| Mutation | ΔG (kcal/mol) | | $\Delta\Delta G_{binding}$ (kcal/mol) | Production run |
|---|---|---|---|---|
| | START-Cer-POPC-Water | START-POPC-Water | START-Cer | (ns) |
| V480A | 2.8 ± 0.3 | 2.7 ± 0.2 | 0.1 ± 0.2 | 150 |

**Table S6. MSλD START-Cer relative binding free energies for the V480A substitution in the presence of a POPC tail in the binding pocket.** The production simulations were run in 5 replicates (30 ns each).

| Protein | Lipid | Adduct | Full Identity Of Lipid | Intensity | Total Carbon Chain Length | Total Carbon Chain Unsaturations | Ion Mode | mz | Mean Of Retention Time | Minimum Of Retention Time | Maximum Of Retention Time |
|---|---|---|---|---|---|---|---|---|---|---|---|
| CERT | Cer(d34:1) | [M+H]+ | [M+H]+: Ceramide (d34:1) =>(d18:1/16:0) | 3000000 | 34 | 1 | pos | 538.5229 | 10.40 | 10.01 | 10.68 |
| CERT | Cer(d*34:1) | [M+Na]+ | [M+Na]+: Ceramide (d*34:1) | 1700000 | 34 | 1 | pos | 560.4967 | 10.43 | 10.05 | 10.62 |
| CERT | Cer(d32:1) | [M+H]+ | [M+H]+: Ceramide (d32:1) =>(d18:1/14:0) | 440000 | 32 | 1 | pos | 510.4906 | 9.10 | 8.46 | 10.70 |
| CERT | Cer(d*32:1) | [M+Na]+ | [M+Na]+: Ceramide (d*32:1) | 340000 | 32 | 1 | pos | 532.4736 | 9.07 | 8.59 | 9.22 |
| CERT | Cer(d34:2) | [M+H]+ | [M+H]+: Ceramide (d34:2) =>(d18:2/16:0) | 260000 | 34 | 2 | pos | 536.5075 | 9.30 | 8.85 | 9.65 |
| CERT | Cer(d*34:2) | [M+Na]+ | [M+Na]+: Ceramide (d*34:2) | 240000 | 34 | 2 | pos | 558.4820 | 9.30 | 8.85 | 9.60 |
| CERT | Cer(d36:1) | [M+H]+ | [M+H]+: Ceramide (d36:1) =>(d18:1/18:0) | 140000 | 36 | 1 | pos | 566.5460 | 11.88 | 11.57 | 12.20 |
| CERT | Cer(d33:1) | [M+H]+ | [M+H]+: Ceramide (d33:1) =>(d17:1/16:0) | 130000 | 33 | 1 | pos | 524.5067 | 9.73 | 9.31 | 10.22 |
| CERT | Cer(d*33:1) | [M+Na]+ | [M+Na]+: Ceramide (d*33:1) | 120000 | 33 | 1 | pos | 546.4909 | 9.82 | 9.27 | 10.59 |
| CERT | Cer(d34:1) | [M+HCOO]- | [M+HCOO]-: Ceramide (d34:1) =>(d18:1/16:0) | 1200000 | 34 | 1 | neg | 582.5033 | 10.42 | 10.14 | 10.60 |
| CERT | Cer(d32:1) | [M+HCOO]- | [M+HCOO]-: Ceramide (d32:1) =>(d18:1/14:0) | 130000 | 32 | 1 | neg | 554.4725 | 9.05 | 8.70 | 9.21 |

**Table S7: Ceramide (dCer) species that were observed in association with CERT by LC-MS/MS.** The different columns describe in order: the lipid transfer protein of interest; the dCer species (d*Cer indicates ceramides that are likely dCer but could also be DHCer), the adduct as which the dCer was observed, and the full identity of the dCer with possible extra information on the fatty acyl composition; the MS intensity by which the dCer was observed; its total carbon chain length and unsaturation; the MS ion mode in which the dCer was observed and at what m/z and LC retention time range.

| Protein | Lipid | Adduct | Full Identity Of Lipid | Intensity | Total Carbon Chain Length | Total Carbon Chain Unsaturations | Ion Mode | mz | Mean Of Retention Time | Minimum Of Retention Time | Maximum Of Retention Time |
|---|---|---|---|---|---|---|---|---|---|---|---|
| GM2A | PC(33:1) | [M+H]+ | [M+H]+: Phosphatidylcholine (33:1) | 470000 | 33 | 1 | pos | 746.5571 | 9.16 | 9.04 | 11.26 |
| GM2A | PC(40:2) | [M+H]+ | [M+H]+: Phosphatidylcholine (40:2) | 420000 | 40 | 2 | pos | 842.6587 | 13.79 | 13.58 | 14.42 |
| GM2A | PC(31:1) | [M+HCOO]- | [M+HCOO]-: Phosphatidylcholine (31:1) => (16:1/15:0) | 92000 | 31 | 1 | neg | 762.5208 | 8.93 | 8.55 | 9.61 |
| LCN1 | PC(32:1) | [M+H]+ | [M+H]+: Phosphatidylcholine (32:1) | 5100000 | 32 | 1 | pos | 732.5501 | 9.34 | 8.58 | 10.19 |
| LCN1 | PC(36:2) | [M+H]+ | [M+H]+: Phosphatidylcholine (36:2) | 5000000 | 36 | 2 | pos | 786.5950 | 10.82 | 9.9 | 11.29 |
| LCN1 | PC(34:1) | [M+H]+ | [M+H]+: Phosphatidylcholine (34:1) | 4900000 | 34 | 1 | pos | 760.5771 | 10.66 | 10.36 | 11.09 |
| LCN1 | PC(34:2) | [M+H]+ | [M+H]+: Phosphatidylcholine (34:2) | 4000000 | 34 | 2 | pos | 758.5638 | 9.59 | 9.19 | 10.61 |
| LCN1 | PC(32:2) | [M+H]+ | [M+H]+: Phosphatidylcholine (32:2) | 2400000 | 32 | 2 | pos | 730.5339 | 8.48 | 8.00 | 9.81 |
| LCN1 | PC(30:1) | [M+H]+ | [M+H]+: Phosphatidylcholine (30:1) | 1800000 | 30 | 1 | pos | 704.5178 | 8.17 | 7.78 | 8.65 |
| LCN1 | PC(33:1) | [M+H]+ | [M+H]+: Phosphatidylcholine (33:1) | 660000 | 33 | 1 | pos | 746.5643 | 10.06 | 6.96 | 10.66 |
| LCN1 | PC(34:3) | [M+H]+ | [M+H]+: Phosphatidylcholine (34:3) | 520000 | 34 | 3 | pos | 756.5482 | 8.70 | 8.30 | 9.44 |
| LCN1 | PC(31:1) | [M+H]+ | [M+H]+: Phosphatidylcholine (31:1) | 440000 | 31 | 1 | pos | 718.5338 | 8.74 | 8.21 | 10.09 |
| LCN1 | PC(36:1) | [M+H]+ | [M+H]+: Phosphatidylcholine (36:1) | 380000 | 36 | 1 | pos | 788.6105 | 12.09 | 11.73 | 12.5 |
| LCN1 | PC(36:4) | [M+H]+ | [M+H]+: Phosphatidylcholine (36:4) | 280000 | 36 | 4 | pos | 782.5635 | 9.53 | 8.43 | 10.43 |
| LCN1 | PC(33:2) | [M+H]+ | [M+H]+: Phosphatidylcholine (33:2) | 260000 | 33 | 2 | pos | 744.5500 | 8.97 | 8.55 | 10.09 |
| LCN1 | PC(38:5) | [M+H]+ | [M+H]+: Phosphatidylcholine (38:5) | 230000 | 38 | 5 | pos | 808.5804 | 9.65 | 9.11 | 10.17 |
| LCN1 | PC(30:0) | [M+H]+ | [M+H]+: Phosphatidylcholine (30:0) | 230000 | 30 | 0 | pos | 706.5332 | 9.13 | 8.68 | 9.45 |
| LCN1 | PC(38:6) | [M+H]+ | [M+H]+: Phosphatidylcholine (38:6) | 210000 | 38 | 6 | pos | 806.5648 | 9.28 | 8.21 | 10.17 |
| LCN1 | PC(32:1) | [M+Na]+ | [M+Na]+: Phosphatidylcholine (32:1) | 150000 | 32 | 1 | pos | 754.5295 | 9.32 | 9.02 | 9.81 |
| LCN1 | PC(34:1) | [M+Na]+ | [M+Na]+: Phosphatidylcholine (34:1) | 140000 | 34 | 1 | pos | 782.5602 | 10.65 | 10.39 | 10.97 |

| | | | | | | | | | | | |
|---|---|---|---|---|---|---|---|---|---|---|---|
| LCN1 | PC(36:2) | [M+Na]+ | [M+Na]+: Phosphatidylcholine (36:2) | 140000 | 36 | 2 | pos | 808.5768 | 10.79 | 10.52 | 11.13 |
| LCN1 | PC(31:0) | [M+H]+ | [M+H]+: Phosphatidylcholine (31:0) | 120000 | 31 | 0 | pos | 720.5490 | 9.51 | 9.19 | 9.89 |
| LCN1 | PC(40:7) | [M+H]+ | [M+H]+: Phosphatidylcholine (40:7) | 120000 | 40 | 7 | pos | 832.5811 | 9.40 | 8.93 | 9.77 |
| LCN1 | PC(34:2) | [M+Na]+ | [M+Na]+: Phosphatidylcholine (34:2) | 110000 | 34 | 2 | pos | 780.5453 | 9.53 | 9.19 | 10.11 |
| LCN1 | PC(30:2) | [M+H]+ | [M+H]+: Phosphatidylcholine (30:2) | 130000 | 30 | 2 | pos | 702.5030 | 7.56 | 7.00 | 8.22 |
| LCN1 | PC(32:1) | [M+HCOO]- | [M+HCOO]-: Phosphatidylcholine (32:1) => (16:1/16:0) | 140000 | 32 | 1 | neg | 776.5365 | 9.33 | 9.03 | 9.79 |
| LCN1 | PC(34:1) | [M+HCOO]- | [M+HCOO]-: Phosphatidylcholine (34:1) => (18:1/16:0) | 110000 | 34 | 1 | neg | 804.5664 | 10.64 | 10.38 | 10.93 |
| LCN1 | PC(34:2) | [M+HCOO]- | [M+HCOO]-: Phosphatidylcholine (34:2) => (16:1/18:1) | 100000 | 34 | 2 | neg | 802.5513 | 9.50 | 9.24 | 10.08 |
| PITPNA | PC(33:1) | [M+H]+ | [M+H]+: Phosphatidylcholine (33:1) | 130000 | 33 | 1 | pos | 746.5789 | 9.68 | 9.14 | 10.54 |
| PITPNA | PC(30:1) | [M+H]+ | [M+H]+: Phosphatidylcholine (30:1) | 120000 | 30 | 1 | pos | 704.5321 | 7.91 | 7.58 | 8.38 |
| PITPNB | PC(32:1) | [M+H]+ | [M+H]+: Phosphatidylcholine (32:1) | 380000 | 32 | 1 | pos | 732.5632 | 9.14 | 7.14 | 10.41 |
| PITPNB | PC(34:1) | [M+H]+ | [M+H]+: Phosphatidylcholine (34:1) | 340000 | 34 | 1 | pos | 760.5938 | 10.4 | 10.07 | 10.9 |
| PITPNB | PC(34:2) | [M+H]+ | [M+H]+: Phosphatidylcholine (34:2) | 210000 | 34 | 2 | pos | 758.5788 | 9.40 | 9.00 | 10.73 |
| PITPNB | PC(30:0) | [M+H]+ | [M+H]+: Phosphatidylcholine (30:0) | 33000 | 30 | 0 | pos | 706.5481 | 8.88 | 8.65 | 9.14 |
| STARD2 | PC(34:1) | [M+H]+ | [M+H]+: Phosphatidylcholine (34:1) | 17000000 | 34 | 1 | pos | 760.5780 | 10.98 | 10.72 | 11.41 |
| STARD2 | PC(36:2) | [M+H]+ | [M+H]+: Phosphatidylcholine (36:2) | 16000000 | 36 | 2 | pos | 786.5944 | 11.13 | 10.76 | 11.64 |
| STARD2 | PC(34:2) | [M+H]+ | [M+H]+: Phosphatidylcholine (34:2) | 11000000 | 34 | 2 | pos | 758.5645 | 9.94 | 9.49 | 10.98 |
| STARD2 | PC(32:1) | [M+H]+ | [M+H]+: Phosphatidylcholine (32:1) | 11000000 | 32 | 1 | pos | 732.5499 | 9.80 | 6.93 | 11.53 |
| STARD2 | PC(32:2) | [M+H]+ | [M+H]+: Phosphatidylcholine (32:2) | 3200000 | 32 | 2 | pos | 730.5345 | 8.74 | 8.34 | 9.50 |
| STARD2 | PC(36:3) | [M+H]+ | [M+H]+: Phosphatidylcholine (36:3) | 2700000 | 36 | 3 | pos | 784.5806 | 10.19 | 9.76 | 11.05 |
| STARD2 | PC(38:6) | [M+H]+ | [M+H]+: Phosphatidylcholine (38:6) | 1700000 | 38 | 6 | pos | 806.5657 | 9.64 | 8.47 | 10.21 |

| | | | | | | | | | | | |
|---|---|---|---|---|---|---|---|---|---|---|---|
| STARD2 | PC(38:5) | [M+H]+ | [M+H]+: Phosphatidylcholine (38:5) | 1700000 | 38 | 5 | pos | 808.5819 | 10.03 | 9.49 | 10.58 |
| STARD2 | PC(40:7) | [M+H]+ | [M+H]+: Phosphatidylcholine (40:7) | 1500000 | 40 | 7 | pos | 832.5814 | 9.79 | 9.49 | 10.12 |
| STARD2 | PC(36:4) | [M+H]+ | [M+H]+: Phosphatidylcholine (36:4) | 1600000 | 36 | 4 | pos | 782.5646 | 9.90 | 8.94 | 10.21 |
| STARD2 | PC(36:1) | [M+H]+ | [M+H]+: Phosphatidylcholine (36:1) | 1300000 | 36 | 1 | pos | 788.6122 | 12.41 | 12.15 | 12.83 |
| STARD2 | PC(30:1) | [M+H]+ | [M+H]+: Phosphatidylcholine (30:1) | 1200000 | 30 | 1 | pos | 704.5179 | 8.54 | 8.10 | 8.91 |
| STARD2 | PC(33:1) | [M+H]+ | [M+H]+: Phosphatidylcholine (33:1) | 1100000 | 33 | 1 | pos | 746.5662 | 10.29 | 9.28 | 10.85 |
| STARD2 | PC(34:3) | [M+H]+ | [M+H]+: Phosphatidylcholine (34:3) | 980000 | 34 | 3 | pos | 756.5484 | 9.09 | 8.60 | 9.96 |
| STARD2 | PC(38:3) | [M+H]+ | [M+H]+: Phosphatidylcholine (38:3) | 670000 | 38 | 3 | pos | 812.6126 | 11.77 | 10.39 | 12.42 |
| STARD2 | PC(38:7) | [M+H]+ | [M+H]+: Phosphatidylcholine (38:7) | 560000 | 38 | 7 | pos | 804.5499 | 8.56 | 8.26 | 9.90 |
| STARD2 | PC(34:1) | [M+Na]+ | [M+Na]+: Phosphatidylcholine (34:1) | 510000 | 34 | 1 | pos | 782.5605 | 10.97 | 10.76 | 11.28 |
| STARD2 | PC(38:4) | [M+H]+ | [M+H]+: Phosphatidylcholine (38:4) | 510000 | 38 | 4 | pos | 810.5961 | 11.27 | 10.81 | 11.59 |
| STARD2 | PC(36:5) | [M+H]+ | [M+H]+: Phosphatidylcholine (36:5) | 500000 | 36 | 5 | pos | 780.5490 | 8.80 | 8.04 | 9.43 |
| STARD2 | PC(40:6) | [M+H]+ | [M+H]+: Phosphatidylcholine (40:6) | 480000 | 40 | 6 | pos | 834.5973 | 10.15 | 9.66 | 10.83 |
| STARD2 | PC(35:1) | [M+H]+ | [M+H]+: Phosphatidylcholine (35:1) | 480000 | 35 | 1 | pos | 774.5956 | 11.55 | 11.08 | 11.91 |
| STARD2 | PC(40:6) | [M+H]+ | [M+H]+: Phosphatidylcholine (40:6) | 450000 | 40 | 6 | pos | 834.5957 | 11.02 | 10.76 | 11.38 |
| STARD2 | PC(36:2) | [M+Na]+ | [M+Na]+: Phosphatidylcholine (36:2) | 450000 | 36 | 2 | pos | 808.5762 | 11.12 | 10.89 | 11.6 |
| STARD2 | PC(34:2) | [M+Na]+ | [M+Na]+: Phosphatidylcholine (34:2) | 330000 | 34 | 2 | pos | 780.5457 | 9.89 | 9.54 | 10.46 |
| STARD2 | PC(31:1) | [M+H]+ | [M+H]+: Phosphatidylcholine (31:1) | 320000 | 31 | 1 | pos | 718.5341 | 9.08 | 8.52 | 9.62 |
| STARD2 | PC(32:1) | [M+Na]+ | [M+Na]+: Phosphatidylcholine (32:1) | 310000 | 32 | 1 | pos | 754.5300 | 9.79 | 9.37 | 10.16 |
| STARD2 | PC(31:0) | [M+H]+ | [M+H]+: Phosphatidylcholine (31:0) | 210000 | 31 | 0 | pos | 720.5496 | 9.82 | 9.58 | 10.21 |
| STARD2 | PC(30:0) | [M+H]+ | [M+H]+: Phosphatidylcholine (30:0) | 210000 | 30 | 0 | pos | 706.5342 | 9.38 | 9.12 | 9.72 |
| STARD2 | PC(36:6) | [M+H]+ | [M+H]+: Phosphatidylcholine (36:6) | 200000 | 36 | 6 | pos | 778.5341 | 8.35 | 7.73 | 9.49 |

| | | | | | | | | | | | |
|---|---|---|---|---|---|---|---|---|---|---|---|
| STARD2 | PC(34:1) | [M+HCOO]- | [M+HCOO]-: Phosphatidylcholine (34:1) => (18:1/16:0) | 540000 | 34 | 1 | neg | 804.5667 | 10.92 | 10.66 | 11.27 |
| STARD2 | PC(36:2) | [M+HCOO]- | [M+HCOO]-: Phosphatidylcholine (36:2) => (18:1/18:1) | 460000 | 36 | 2 | neg | 830.5823 | 11.09 | 10.8 | 11.56 |
| STARD2 | PC(32:1) | [M+HCOO]- | [M+HCOO]-: Phosphatidylcholine (32:1) => (16:1/16:0) | 410000 | 32 | 1 | neg | 776.5366 | 9.61 | 9.28 | 10.13 |
| STARD2 | PC(34:2) | [M+HCOO]- | [M+HCOO]-: Phosphatidylcholine (34:2) => (18:1/16:1) | 380000 | 34 | 2 | neg | 802.5512 | 9.80 | 9.48 | 10.38 |
| STARD2 | PC(32:2) | [M+HCOO]- | [M+HCOO]-: Phosphatidylcholine (32:2) => (16:1/16:1) | 120000 | 32 | 2 | neg | 774.5207 | 8.69 | 8.28 | 9.32 |
| STARD2 | PC(34:1) | [M+HCOO+1x (HCOONa)]- | [M+HCOO+1x(HCOONa)]-: Phosphatidylcholine (34:1) | 110000 | 34 | 1 | neg | 872.5539 | 10.92 | 10.67 | 11.22 |
| STARD2 | PC(36:2) | [M+HCOO+1x (HCOONa)]- | [M+HCOO+1x(HCOONa)]-: Phosphatidylcholine (36:2) | 90000 | 36 | 2 | neg | 898.5689 | 11.07 | 10.8 | 11.52 |
| STARD2 | PC(34:1) | [M+HCOO+2x (HCOONa)]- | [M+HCOO+2x(HCOONa)]-: Phosphatidylcholine (34:1) | 87000 | 34 | 1 | neg | 940.5403 | 10.92 | 10.66 | 11.22 |
| STARD2 | PC(32:1) | [M+HCOO+1x (HCOONa)]- | [M+HCOO+1x(HCOONa)]-: Phosphatidylcholine (32:1) | 76000 | 32 | 1 | neg | 844.5222 | 9.68 | 9.32 | 10.04 |
| STARD2 | PC(34:2) | [M+HCOO+1x (HCOONa)]- | [M+HCOO+1x(HCOONa)]-: Phosphatidylcholine (34:2) | 76000 | 34 | 2 | neg | 870.5377 | 9.80 | 9.53 | 10.3 |
| STARD2 | PC(34:1) | [M+HCOO+3x (HCOONa)]- | [M+HCOO+3x(HCOONa)]-: Phosphatidylcholine (34:1) | 62000 | 34 | 1 | neg | 1008.5268 | 10.92 | 10.67 | 11.17 |
| STARD10 | PC(36:2) | [M+H]+ | [M+H]+: Phosphatidylcholine (36:2) | 11000000 | 36 | 2 | pos | 786.5987 | 10.68 | 10.29 | 11.04 |
| STARD10 | PC(34:2) | [M+H]+ | [M+H]+: Phosphatidylcholine (34:2) | 8700000 | 34 | 2 | pos | 758.5674 | 9.42 | 9.00 | 9.90 |
| STARD10 | PC(34:1) | [M+H]+ | [M+H]+: Phosphatidylcholine (34:1) | 4400000 | 34 | 1 | pos | 760.5822 | 10.53 | 10.2 | 10.85 |
| STARD10 | PC(32:1) | [M+H]+ | [M+H]+: Phosphatidylcholine (32:1) | 3800000 | 32 | 1 | pos | 732.5533 | 9.25 | 7.41 | 11.63 |
| STARD10 | PC(32:2) | [M+H]+ | [M+H]+: Phosphatidylcholine (32:2) | 1700000 | 32 | 2 | pos | 730.5380 | 8.29 | 7.83 | 9.48 |
| STARD10 | PC(36:3) | [M+H]+ | [M+H]+: Phosphatidylcholine (36:3) | 1600000 | 36 | 3 | pos | 784.5838 | 9.75 | 9.33 | 10.22 |
| STARD10 | PC(30:1) | [M+H]+ | [M+H]+: Phosphatidylcholine (30:1) | 720000 | 30 | 1 | pos | 704.5206 | 8.07 | 7.12 | 10.44 |
| STARD10 | PC(34:3) | [M+H]+ | [M+H]+: Phosphatidylcholine (34:3) | 680000 | 34 | 3 | pos | 756.5517 | 8.59 | 8.13 | 9.28 |
| STARD10 | PC(36:4) | [M+H]+ | [M+H]+: Phosphatidylcholine (36:4) | 570000 | 36 | 4 | pos | 782.5677 | 8.83 | 8.38 | 9.73 |

| | | | | | | | | | | | |
|---|---|---|---|---|---|---|---|---|---|---|---|
| STARD10 | PC(35:2) | [M+H]+ | [M+H]+: Phosphatidylcholine (35:2) | 450000 | 35 | 2 | pos | 772.5833 | 10.02 | 9.62 | 10.43 |
| STARD10 | PC(38:6) | [M+H]+ | [M+H]+: Phosphatidylcholine (38:6) | 400000 | 38 | 6 | pos | 806.5692 | 9.25 | 8.08 | 10.01 |
| STARD10 | PC(36:2) | [M+Na]+ | [M+Na]+: Phosphatidylcholine (36:2) | 350000 | 36 | 2 | pos | 808.5821 | 10.68 | 10.33 | 11.00 |
| STARD10 | PC(40:7) | [M+H]+ | [M+H]+: Phosphatidylcholine (40:7) | 320000 | 40 | 7 | pos | 832.5849 | 9.36 | 8.96 | 9.64 |
| STARD10 | PC(38:4) | [M+H]+ | [M+H]+: Phosphatidylcholine (38:4) | 290000 | 38 | 4 | pos | 810.6002 | 10.11 | 9.54 | 10.75 |
| STARD10 | PC(38:5) | [M+H]+ | [M+H]+: Phosphatidylcholine (38:5) | 270000 | 38 | 5 | pos | 808.5844 | 9.62 | 9.26 | 9.97 |
| STARD10 | PC(33:2) | [M+H]+ | [M+H]+: Phosphatidylcholine (33:2) | 190000 | 33 | 2 | pos | 744.5533 | 8.82 | 8.42 | 9.32 |
| STARD10 | PC(34:2) | [M+Na]+ | [M+Na]+: Phosphatidylcholine (34:2) | 170000 | 34 | 2 | pos | 780.5494 | 9.42 | 9.04 | 9.85 |
| STARD10 | PC(38:2) | [M+H]+ | [M+H]+: Phosphatidylcholine (38:2) | 160000 | 38 | 2 | pos | 814.6322 | 11.99 | 11.6 | 12.41 |
| STARD10 | PC(38:7) | [M+H]+ | [M+H]+: Phosphatidylcholine (38:7) | 140000 | 38 | 7 | pos | 804.5533 | 8.21 | 7.83 | 8.58 |
| STARD10 | PC(36:2) | [M+HCOO]- | [M+HCOO]-: Phosphatidylcholine (36:2) => (18:1/18:1) | 270000 | 36 | 2 | neg | 830.5859 | 10.72 | 10.48 | 11.00 |
| STARD10 | PC(34:1) | [M+HCOO]- | [M+HCOO]-: Phosphatidylcholine (34:1) => (18:1/16:0) | 100000 | 34 | 1 | neg | 804.5701 | 10.57 | 10.35 | 10.86 |
| CERT | PC(34:1) | [M+H]+ | [M+H]+: Phosphatidylcholine (34:1) | 610000 | 34 | 1 | pos | 760.5778 | 10.31 | 9.97 | 11.14 |
| CERT | PC(32:1) | [M+H]+ | [M+H]+: Phosphatidylcholine (32:1) | 250000 | 32 | 1 | pos | 732.5490 | 9.03 | 6.24 | 10.66 |

**Table S8: Phosphatidylcholine (PC) species observed with CERT and other known PC-binding lipid transfer proteins by LC-MS/MS.** The different columns describe in order: the lipid transfer protein of interest; the PC species, the adduct as which the PC was observed, and the full identity of the PC with possible extra information on the fatty acyl composition; the MS intensity by which the PC was observed; its total carbon chain length and unsaturation; the MS ion mode in which the PC was observed and at what m/z and LC retention time range.

| Residue/lipid | Candidate atoms |
|---|---|
| **ALA** | CA HA CB HB1 HB2 HB3 |
| **ARG** | CA HA CB HB1 HB2 CG HG1 HG2 |
| **ASN** | CA HA CB HB1 HB2 |
| **ASP** | CA HA CB HB1 HB2 |
| **CYS** | CA HA CB HB1 HB2 |
| **GLN** | CA HA CB HB1 HB2 CG HG1 HG2 |
| **GLU** | CA HA CB HB1 HB2 CG HG1 HG2 |
| **GLY** | CA HA1 HA2 |
| **HSD** | CA HA CB HB1 HB2 |
| **HSE** | CA HA CB HB1 HB2 |
| **HSP** | CA HA CB HB1 HB2 CG |
| **ILE** | CA HA CB HB CG1 HG11 HG12 CG2 HG21 HG22 HG23 CD HD1 HD2 HD3 |
| **LEU** | CA HA CB HB1 HB2 CG HG CD1 HD11 HD12 HD13 CD2 HD21 HD22 HD23 |
| **LYS** | CA HA CB HB1 HB2 CG HG1 HG2 CD HD1 HD2 CE HE1 HE2 |
| **MET** | CA HA CB HB1 HB2 CG HG1 HG2 CE HE1 HE2 HE3 |
| **PHE** | CA HA CB HB1 HB2 CG CD1 HD1 CD2 HD2 CE1 HE1 CE2 HE2 CZ HZ |
| **PRO** | CA HA CB HB1 HB2 CD HD1 HD2 CG HG1 HG2 |
| **SER** | CA HA CB HB1 HB2 |
| **THR** | CA HA CB HB CG2 HG21 HG22 HG23 |
| **TRP** | CA HA CB HB1 HB2 CG CD1 HD1 CD2 CE3 HE3 CZ3 HZ3 CH2 HH2 CZ2 HZ2 |
| **TYR** | CA HA CB HB1 HB2 CG CD1 HD1 CD2 HD2 CE1 HE1 |
| **VAL** | CA HA CB HB CG1 HG11 HG12 HG13 CG2 FG21 HG22 HG23 |
| **POPC/POPE/ POPI/POPS/ POPA/POGL** | C23 H3R H3S C24 H4R H4S C25 H5R H5S C26 H6R H6S C27 H7R H7S C28 H8R H8S C29 H9S1 C210 H10S1 C211 H11R H11S C212 H12R H12S C213 H13R H13S C214 H14R H14S C215 H15R H15S C216 H16R H16S C217 H17R H17S C218 H18R H18S H18T C33 H3X H3Y C34 H4X H4Y C35 H5X H5Y C36 H6X H6Y C37 H7X H7Y C38 H8X H8Y C39 H9X H9Y C310 H10X H10Y C311 H11X H11Y C312 H12X H12Y C313 H13X H13Y C314 H14X H14Y C315 H15X H15Y C316 H16X H16Y H16Z |
| **CER180** | C5S C6S H6S H6T C7S H7S H7T C8S H8S H8T C9S H9S H9T C10S H10S H10T C11S H11S H11T C12S H12S H12T C13S H13S H13T C14S H14S H14T C15S H15S H15T C16S H16S H16T C17S H17S H17T C18S H18S H18T H18U C2F H2F H2G C3F H3F H3G C4F H4F H4G C5F H5F H5G C6F H6F H6G C7F H7F H7G C8F H8F H8G C9F H9F H9G C10F H10F H10G C11F H11F H11G C12F H12F H12G C13F H13F H13G C14F H14F H14G C15F H15F H15G C16F H16F H16G C17F H17F H17G C18F H18F H18G H18H |
| **CHOL** | C4 H4A H4B C5 C6 H6 C7 H7A H7B C8 H8 C14 H14 C15 H15A H15B C16 H16A H16B C17 H17 C13 C18 H18A H18B H18C C12 H12A H12B C11 H11A H11B C9 H9 C10 C19 H19A H19B H19C C1 H1A H1B C2 H2A H2B C20 H20 C21 H21A H21B H21C C22 H22A H22B C23 H23A H23B C24 H24A H24B C25 H25 C26 H26A H26B H26C C27 H27A H27B H27C |

**Table S9. Candidate atoms for the hydrophobic interaction analysis.** The atom names correspond to the nomenclature of the CHARMM36 force field.

**Movie S1.** POPC lipid inserts a tail in the cavity in the simulation of the apo+ER_r1. Simulation excerpt is from 400-950ns. POPC lipid is shown in purple. Ω1 loop colored orange, Ω4 loop colored cyan and START domain colored grey. Membrane lipid molecules are shown in licorice representation and for the sake of clarity, water molecules are not shown and only a subset of the lipids are shown.

**Movie S2A.** POPC inserts a tail in the cavity in the simulation of the holo+Neu_r1. Simulation excerpt is from 800-950ns. Ceramide lipid is shown in yellow, POPC lipid is shown in purple. Ω1 loop colored orange, Ω4 loop colored cyan and START domain colored grey. Membrane lipid molecules are shown in licorice representation and for the sake of clarity, water molecules are not shown and only a subset of the lipids are shown.

**Movie S2B.** Release of the bound ceramide from the cavity into the bilayer facilitated by presence of POPC tail in the cavity in the simulation of the holo+Neu_r1. Simulation excerpt is from 1380-1550ns. Ceramide lipid is shown in yellow, POPC lipid is shown in purple. Ω1 loop colored orange, Ω4 loop colored cyan and START domain colored grey. Membrane lipid molecules are shown in licorice representation and for the sake of clarity, water molecules are not shown and only a subset of the lipids are shown.

**Movie S2C.** Released ceramide returns to the cavity when POPC tails exits the cavity in the simulation of the holo+Neu_r1. Simulation excerpt is from 1570-1770ns. Ceramide lipid is shown in yellow, POPC lipid is shown in purple. Ω1 loop colored orange, Ω4 loop colored cyan and START domain colored grey. Membrane lipid molecules are shown in licorice representation and for the sake of clarity, water molecules are not shown and only a subset of the lipids are shown.

**Movie S3.** POPC inserts a tail in the cavity in the simulation of the holo+Golgi_r1. Simulation excerpt is from 810-940ns. Ceramide lipid is shown in yellow, POPC lipid is shown in purple. Ω1 loop colored orange, Ω4 loop colored cyan and START domain colored grey. Membrane lipid molecules are shown in licorice representation and for the sake of clarity, water molecules are not shown and only a subset of the lipids are shown.

**Movie S4.** Release of the ceramide from the cavity followed by uptake of POGL into the cavity in the simulation of the Holo+Golgi+POPC_r1. Simulation excerpt is from 100-1500ns. Ceramide lipid is shown in yellow, POPC lipid is shown in purple, POGL lipid is shown in magenta. Ω1 loop colored orange, Ω4 loop colored cyan and START domain colored grey. Membrane lipid molecules are shown in licorice representation and for the sake of clarity, water molecules are not shown and only a subset of the lipids are shown.
